# The prion-like domain of the chloroplast RNA binding protein CP29A is required for cold-induced phase separation next to nucleoids and supports RNA splicing and translation during cold acclimation

**DOI:** 10.1101/2023.09.29.560215

**Authors:** Julia Legen, Benjamin Lenzen, Nitin Kachariya, Stephanie Feltgen, Yang Gao, Simon Mergenthal, Willi Weber, Enrico Klotzsch, Reimo Zoschke, Michael Sattler, Christian Schmitz-Linneweber

## Abstract

*Arabidopsis thaliana* is capable of producing photosynthetic tissue with active chloroplasts at temperatures as low as 4°C, and this process depends on the presence of the nuclear-encoded, chloroplast-localized RNA-binding protein CP29A. In this study, we demonstrate that CP29A undergoes phase separation in vitro and in vivo in a temperature-dependent manner, which is mediated by a prion-like domain (PLD) located between the two RNA recognition motif (RRM) domains of CP29A. The resulting droplets display liquid-like properties and are found in close proximity to chloroplast nucleoids. The PLD is required to support chloroplast RNA splicing and translation in cold-treated tissue. Together, our findings suggest that plant chloroplast gene expression is compartmentalized by inducible condensation of CP29A at low temperatures, a mechanism that could play a crucial role for plant cold resistance.

## Introduction

Phase separation (PS) is a phenomenon that can lead to the formation of membraneless compartments within cells (Shin and Brangwynne, 2017). These compartments are a dynamic alternative to organelles, as they lack a physical, membranous boundary and can be modulated by changes in protein concentrations, metabolites, salt or temperature (Alberti and Hyman, 2021). Compartments induced by PS are important for efficiently organizing cellular processes by concentrating specific factors in their appropriate place and at the right time. Unlike irreversible cellular aggregation, these compartments are dynamically formed and dissolved and have a viscoelastic-dynamic, fluid nature, which makes them flexible and adaptable. PS has been demonstrated to be crucial for rapid responses to changing environments, and can be induced by shifts in temperature and cellular homeostasis (Nott et al., 2015; Dignon et al., 2019). Intrinsically disordered protein domains (IDRs) are often at the heart of PS phenomena, driving the formation of cellular membrane-less compartments through multivalent interactions (Kato et al., 2012; Forman-Kay and Mittag, 2013). RNA can act as a seed and essential component for phase-separated compartments, with RNA-binding proteins being over-represented among factors described to phase-separate (Roden and Gladfelter, 2021). PS and the resulting RNA granules play a crucial role in a number of processes involving RNA binding proteins, including ribosome maturation, RNA splicing, ribonucleoprotein biogenesis, and RNA degradation.

In plants, PS phenomena are just beginning to be characterized (Emenecker et al., 2020; Gutierrez-Beltran et al., 2023; Kang and Xu, 2023), but it is already clear that condensates contribute to stress resilience and signaling (Field et al., 2023; Solis-Miranda et al., 2023; Kearly et al., 2022). Almost all of the PS events described in land plants occur in the cytosol or the cytoplasm (Jung et al., 2020; Xie et al., 2021; Fang et al., 2019; Shang et al., 2023; Kim et al., 2021; Tan et al., 2023; Zavaliev et al., 2020; Zhang et al., 2022; Huang et al., 2021b; Hoffmann et al., 2023; Wang et al., 2023; Ruiz-Solaní et al., 2023), but there is one notable exception: In Arabidopsis chloroplasts, an ankyrin protein has been shown to undergo phase separation to facilitate sorting and transport of proteins into the thylakoids (Ouyang et al., 2020). Phase separation also plays a role in chloroplasts of green algae, where the chloroplast pyrenoid, a structure required for carbon concentration around RuBisCo, has been shown to assemble via condensation (Freeman Rosenzweig et al., 2017). There are also reports of RNA granules in the chloroplast of a green alga, but whether they are based on PS phenomena is unclear (Uniacke and Zerges, 2009, 2008). Similarly, heat inducible stress granules were observed in Arabidopsis chloroplasts, but the influence of PS is unclear (Chodasiewicz et al., 2020).

The chloroplast RNA-binding protein CP29A is a member of the cpRNP protein family, which has eleven representatives in Arabidopsis, all localizing to chloroplasts (Ruwe et al., 2011). Unlike the other 10 family members, CP29A’s two RNA recognition motifs (RRMs) are spaced by a linker of 84 amino acids. CP29A responds to different external and internal stimuli, accumulating to higher levels in the cold and being repressed by other stressors and abscisic acid (Kupsch et al., 2012; Raab et al., 2006). CP29A associates with a large number of chloroplast transcripts (Kupsch et al., 2012). A homologue of CP29A in tobacco is required for stabilizing chloroplast RNAs in vitro (Nakamura et al., 2001). Null mutants of CP29A exhibit a pale center of the leaf rosette after long-term exposure to low temperatures, with defects in chloroplast RNA processing and accumulation (Kupsch et al., 2012). When grown at regular growth temperatures, CP29A-deficient plants do not show RNA defects or macroscopic changes relative to wt plants, indicating that CP29A has evolved to support cold acclimation in *Arabidopsis thaliana* (Kupsch et al., 2012). The temperature range in which CP29A is relevant does not fall within the realm of stress, but rather within a range that plants such as *A. thaliana* are constantly exposed to in temperate zones, specifically from 4°C to 12°C. Therefore, CP29A is a factor that is crucial for the daily acclimatization to low temperatures, which are to be expected throughout the year in temperate regions. The molecular mechanisms by which CP29A achieves this acclimatization of plastidic RNA metabolism and prevents bleaching remains unclear, nor was it determined whether the effects on RNA maturation and accumulation observed in pale tissue were of primary origin or are pleiotropic effects of the non-photosynthetic tissue. We report here that CP29A is capable of forming liquid-like droplets in vitro and in vivo via its prion-like domain (PLD) and that the PLD is required for chloroplast splicing, translation and cold resistance.

## Results

### The organellar RNA-binding protein CP29A has a prion-like domain, which is required for phase separation in vitro

The protein CP29A contains two RNA recognition motifs (RRMs) that are separated by a linker domain consisting of 84 amino acids. This linker has a repetitive, low complexity sequence with charged and polar residues, and is intrinsically disordered with a prion-like amino acid composition according to *in silico predictions* using the PLAAC and the SPOTD tool (Figure 1A; Lancaster et al., 2014; Hanson et al., 2019). To determine whether CP29A can undergo phase separation, we used recombinant, purified CP29A and monitored its behavior at different temperatures. We found that the CP29A solution becomes turbid at low - but still physiological - temperatures within a few minutes but returns to full translucence when brought back to room temperature (Figure 1B). By contrast, the protein precipitates irreversibly when heated to 37°C (Figure 1B). Next, we analyzed whether full-length CP29A, the PLD alone or a PLD-deletion protein can form liquid droplets. For the PLD deletion, we removed 60 amino acids comprising the glycine-rich region, but still leaving a linker between the RRMs of 16 amino acids (Figure S1A). With the exception of CP29A, all other cpRNP family members have short linkers; for example CP28A has a linker of 18 residues and CP33B has 26 residues (Figure S1A). We reasoned that within this protein family, the two RRM domains are functional if distanced by a short linker, although we cannot exclude structural effects of the PLD deletion on the RRM domains. For visualization, we fused the different protein versions N-terminally to mNeongreen, while omitting the organellar-targeting sequence, since it is cleaved off in vivo. We expressed the protein variants in *E. coli* and purified them (Figure S1B). Indeed, mNeongreen::CP29A formed spherical liquid droplets and wetted the glass surface after settling to the bottom of the dish (Figure 1C; Movie S1). Droplet size increased over time and droplets started to coalesce, which is indicative of the liquid nature of the phase being formed (Figure 1D). Similarly, the PLD alone assembled droplets that fused, while the PLD-less CP29A protein formed irregularly shaped aggregates that did not show fusion (Figure 1D). To investigate the dynamics of mNeongreen::CP29A within droplets, we performed fluorescence recovery after photobleaching (FRAP) experiments. After high-intensity laser irradiation, fluorescence recovers with a halftime of 3.9±0.2 sec for the full length protein, while PLD alone (5.3±0.4 sec) and CP29AΔPLD (9.9±0.7 sec) recovery is slower (Figure 1E). Furthermore, CP29AΔPLD aggregates did not show full recovery after photobleaching, but plateaued at 69 percent demonstrating low mobility of proteins within this phase, while both full length and PLD CP29A showed a recovery of greater than 95 percent (Figure 1E).

**Figure 1:**
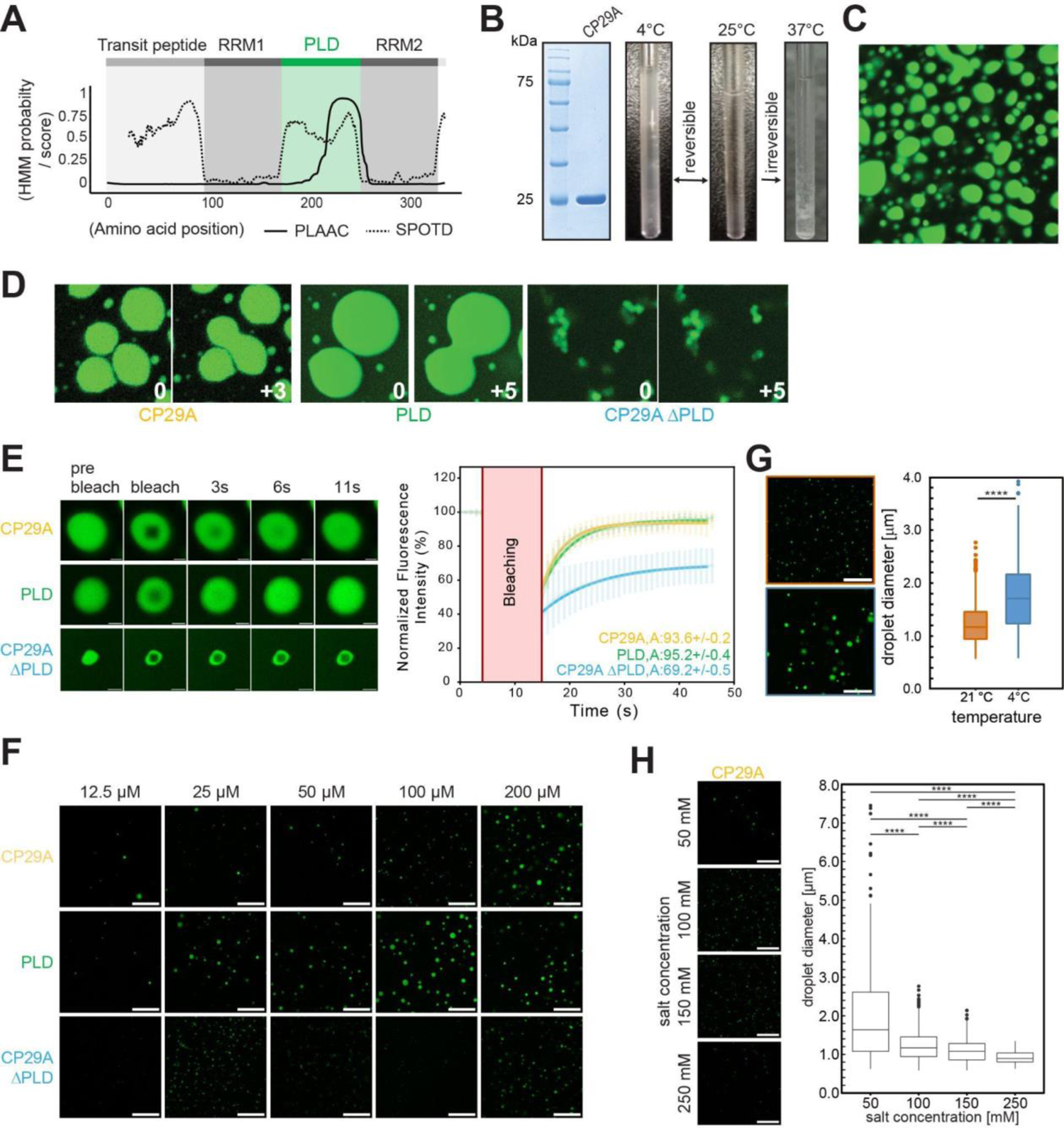
CP29A forms condensates in vitro (A) Schematic diagram of the CP29A protein with its two RRM domains. Below, results of disordered domain predictions within CP29A using the PLAAC and SpotD tools are shown. Scores for both predictors suggest disorderness in the linker between the two RRM domains, including a predicted prion-like domain. (B) Left: Coomassie stain of purified recombinant CP29A excluding the signal peptide, separated by SDS-PAGE. Right: Glass vial with 80 µM CP29A solution is shifted between different temperatures to demonstrate reversible turbidity changes. (C) In vitro phase-separation assay of 100 nM mNG::CP29A with 10% (w/v) PEG4000 treatment for 1 min prior to imaging; imaged with confocal microscopy. (D) Fusion events of mNG::CP29A and mNG:::PLD droplets; conditions as in (C); numbers indicate seconds passed between frames shown; note that CP29AΔPLD forms aggregates that were never observed to fuse. (E) Quantification of fluorescent recovery after photobleaching (FRAP) kinetics with mNG::CP29A, CP29AΔPLD and mNGPLD; all at 100 nM. Error bars = SD; Data are means of the fluorescence values from at least 10 experiments per protein. The images were taken every second over 40 to 50 s to document fluorescence recovery, each time point was normalized to the value before photobleaching. Note that the full-length protein and the PLD recover at a very similar rate, while the PLD-less protein recovers much more slowly. A = plateau intensity as an indicator of the mobile protein fraction. Scale bar: 2 µm (F) Phase separation of mNG::CP29A, mNG:::PLD droplets and CP29AΔPLD at different protein concentrations, scale bar: 20 µm. The experiment was performed three times; representative results are shown. (G) Microscopy of mnG::CP29A droplets (left) and their size distribution (right) of mNG::CP29A at 21°C and 4°C, respectively (50 µM protein; 100 mM NaCl), scale bar 20 µm. (H) Microscopy of phase separation (left) and size distribution (right) of mNG::CP29A at different salt concentrations, scale bar: 20 µm.

As anticipated, our results revealed a positive correlation between droplet formation and protein concentration, as shown in Figure 1F for both the full-length and PLD-only proteins. The PLD-only protein resulted in more droplets than the full-length protein at lower protein concentrations (25 µM), while the PLD-less CP29A did show aggregates at all concentrations assayed (Figure 1F). We also exposed the full-length protein to low temperatures and observed that droplet size increased significantly in the cold (Figure 1G), with an average diameter of 1.2 μm (±0.4 nm) at 21°C and 1.8 (±0.7) μm at 4°C (n = 1497 and n = 197 droplets at 21°C and 4°C, respectively). We next tested the impact of salt on droplet formation and found that concentrations of 100 mM and 150 mM facilitated droplet formation, while droplet-size decreased at higher salt concentrations (Figure 1H). The negative impact of higher salt concentrations on the full-length protein is also observed at low temperatures in turbidity assays (Figure S1C) and can be interpreted as a suppression of intermolecular interactions by electrostatic shielding. Taken together, these data established that CP29A undergoes phase separation in vitro to form liquid droplets, which depends on its PLD and is supported by low temperatures.

### NMR analysis of CP29A indicates molecular interactions of the PLD within the condensed phase

Next, we utilized nuclear magnetic resonance (NMR) spectroscopy along with other biophysical methods in order to investigate the conformation of CP29A at residue level. We expressed and purified the individual RRM domains and the PLD linker and assigned the NMR backbone chemical shifts. Our analysis revealed that CP29A harbors two canonical RRM domains with β_1_-ɑ_1_-β_2_-β_3_-ɑ_2_-β_4_-β_5_ and β_1_-ɑ_1_-β_2_-β_3_-ɑ_2_-β_4_ secondary structure for RRM1 and RRM2, respectively. Furthermore, NMR chemical shifts indicate that the PLD linker connecting the two RRMs is intrinsically disordered (Figure 2A and Figure S1B, D, E). Notably, the PLD linker of CP29A comprises low-complexity regions with repeats of serine (17%), arginine (9%), tyrosine (7%), and glutamate (6%; Figure 2B, Figure S1A), which can potentially engage in electrostatic and cation-π interactions and is rich in glycines (34%; Figure 2B). These features can mediate interactions in phase-separated condensates (Wang et al., 2018b; Qamar et al., 2018). To study the phase separation potential of CP29A in detail, we monitored changes of full length CP29A at temperatures ranging from 298K to 275K in NMR fingerprint spectra. Extensive line-broadening of amide signals was observed at lower temperatures, consistent with reduced molecular tumbling of the protein in a disperse phase (Figure 2C, D; Figure 1B). The extent of line-broadening was significantly larger than expected from the temperature dependence of the molecular tumbling, suggesting that it is due to molecular interactions in the condensed phase.

**Figure 2:**
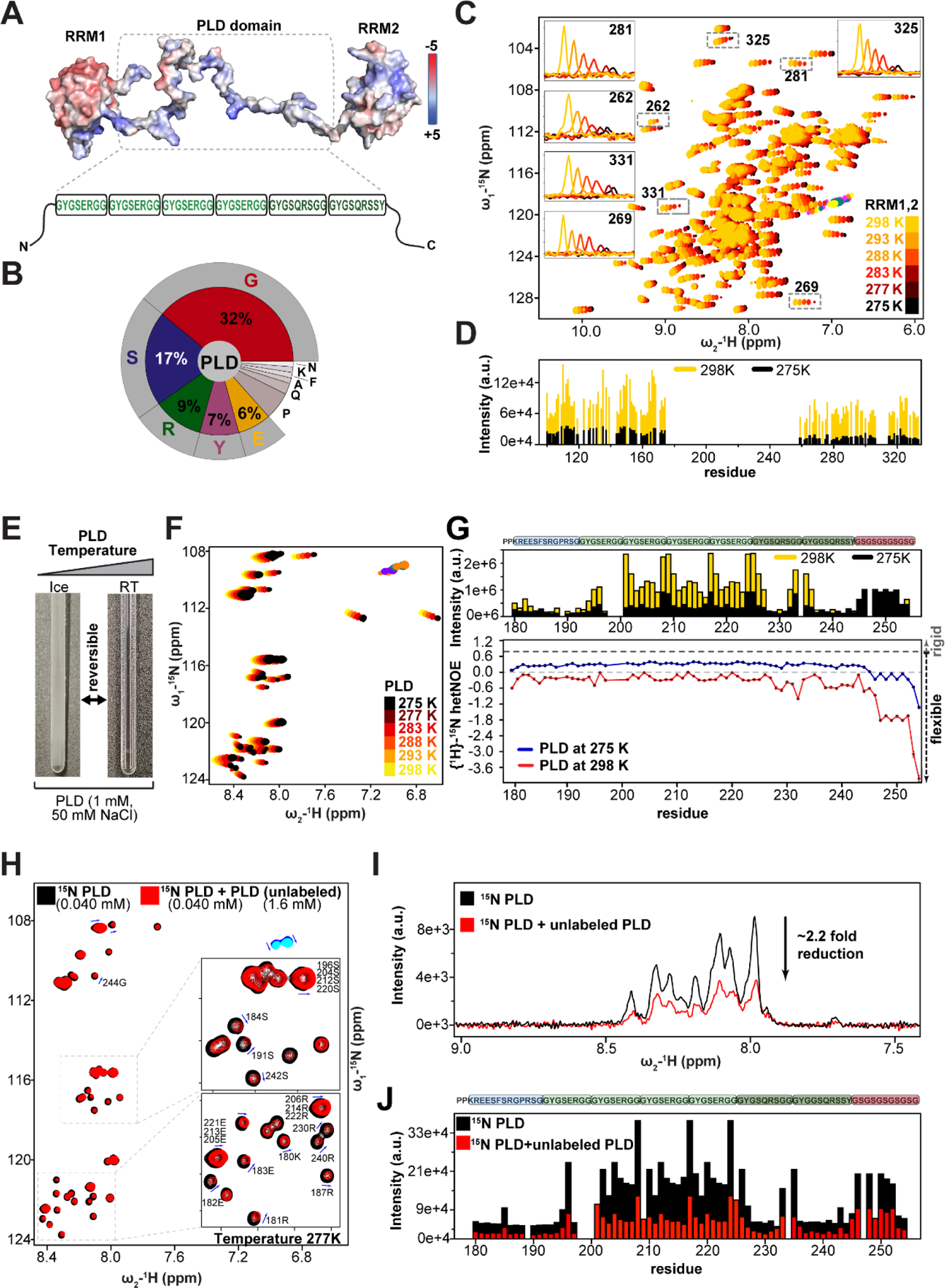
Temperature dependent biophysical characterization of tandem RRMs and the PLD linker of CP29A. (A) Structural model of tandem RRMs of CP29A with electrostatic surface potential rendering (positive (blue) to negative (red) in [kJ/mol/e] as indicated) and the primary sequence of its PLD linker. Repeated sequence elements are boxed. (B) Pie-chart shown for the PLD domain with the proportion of each amino acid. (C) Overlay of ^1^H, ^15^N HSQC spectra of 90 µM concentration of tandem RRMs of CP29A measured at different temperatures. The inset shows the line-broadening with 1D slices of the cross peaks in the 2D experiment upon temperature shift from 298K to 275K. (D) Boxplot for signal intensities of amide protons for spectra shown in figure (C) at 298K (yellow) and at 275K (black). (E) Turbidity changes of a 50 mM PLD solution upon change from ice to room temperature. (F) Overlay of a series of ^1^H, ^15^N HSQC spectra of the PLD linker with increasing temperature from 275K to 298K. Line-broadening of NMR signals is observed at 275K and 277K. (G) Upper panel: Box plot of the intensity of amide proton signals in (F) at 298K and 275K, in yellow and black color, respectively. Lower panel: Comparison of {^1^H}-^15^N heteronuclear NOE of the PLD at 298K and 277K shown in red and blue, respectively. (H) Overlay spectra of the ^15^N labeled PLD alone (black) and in presence of 40-fold excess of unlabeled PLD (red) measured at 277K. Blue arrows indicate smallchemical shift changes. (I) Overlay of 1D amide proton signals for the samples and conditions shown in figure (H). Intensity reduction of the signals is indicated with an arrow. (J) Box plot for signal intensities in the spectra shown in figure (H).

Since the PLD linker is responsible for phase separation of CP29A (Figure 1), we purified the PLD alone and analyzed its biophysical properties (Figure 2E; Figure S1B, D middle panel). By lowering the temperature, noticeable line-broadening of amide signals was observed, similar to the full-length protein, consistent with reduced mobility and condensate formation in disperse solution (Figure 2F, G upper panel). To investigate the conformational dynamics of the PLD in solution, {^1^H}-^15^N heteronuclear nuclear Overhauser effect (hetNOE) experiments were performed at 298K and 275K. These data demonstrate that the PLD has overall reduced backbone flexibility at sub-nanosecond time scales at lower temperature with pronounced rigidification for glycines and serines between residues 245-254 (Figure 2G lower panel). The overall flexibility of the PLD is also seen in a Kratky plot derived from small-angle X-ray scattering (SAXS; Figure S2A). The pairwise-distance distribution suggests a broad range of conformations of the protein up to 100 Å pairwise distances, indicating that the PLD dynamically samples a wide range of conformations in solution (Figure S2B, C; Table S1). In summary, the low complexity region of the PLD of CP29A is unstructured, flexible and undergoes temperature-dependent phase separation similar to previously characterized known PLDs (Franzmann and Alberti, 2019; Wang et al., 2018b).

To confirm that the observed NMR line-broadening reflects reduced tumbling in the condensed phase based on intermolecular interactions, we conducted an NMR titration of ^15^N-labeled PLD with unlabeled PLD at lower temperature (Figure 2H, I). Changes in ^15^N amide chemical shifts and line-broadening, reflected by an 2.2-fold intensity reduction.2 upon the addition of 40-fold molar excess of unlabeled PLD protein, indicated that the ^15^N-labeled protein participates in phase separation with the unlabeled protein in a condensed phase. The most prominent NMR spectral changes are observed for the Gly-Tyr-Gly-Ser-Glu-Arg-Gly-Gly repeats, consistent with weak electrostatic intermolecular interactions (Figure 2J). In sum, our NMR analysis demonstrates high flexibility of the PLD in free solution, which is reduced at lower temperatures consistent with charged interactions that contribute to condensed phase formation.

### CP29A localizes to granules proximal to nucleoids in vivo

We next investigated the potential of CP29A to undergo phase separation within plant chloroplasts. We generated plant lines that express mGFP fused to full-length CP29A (mGFP::CP29Afl), and CP29A without its PLD (mGFP::CP29AΔPLD), both in a *cp29a* null mutant background. The expression of both constructs was driven by the native CP29A promoter, and their expression levels were comparable (Figure S3A). Protoplasts from the mGFP::CP29Afl lines exhibited punctate signals when analyzed by confocal microscopy, which increased in number over time when incubated at 8°C, while protoplasts from the mGFP::CP29AΔPLD plant lines showed much fewer granular signals (Figure 3A). These findings suggest that the PLD of CP29A is strongly supportive of efficient droplet formation in vivo, but also shows that recruitment into granular structures is still possible for the truncated CP29A protein albeit at much lower efficiency. Furthermore, our data indicate that low temperatures not only promote droplet formation in vitro but also in living cells.

**Figure 3:**
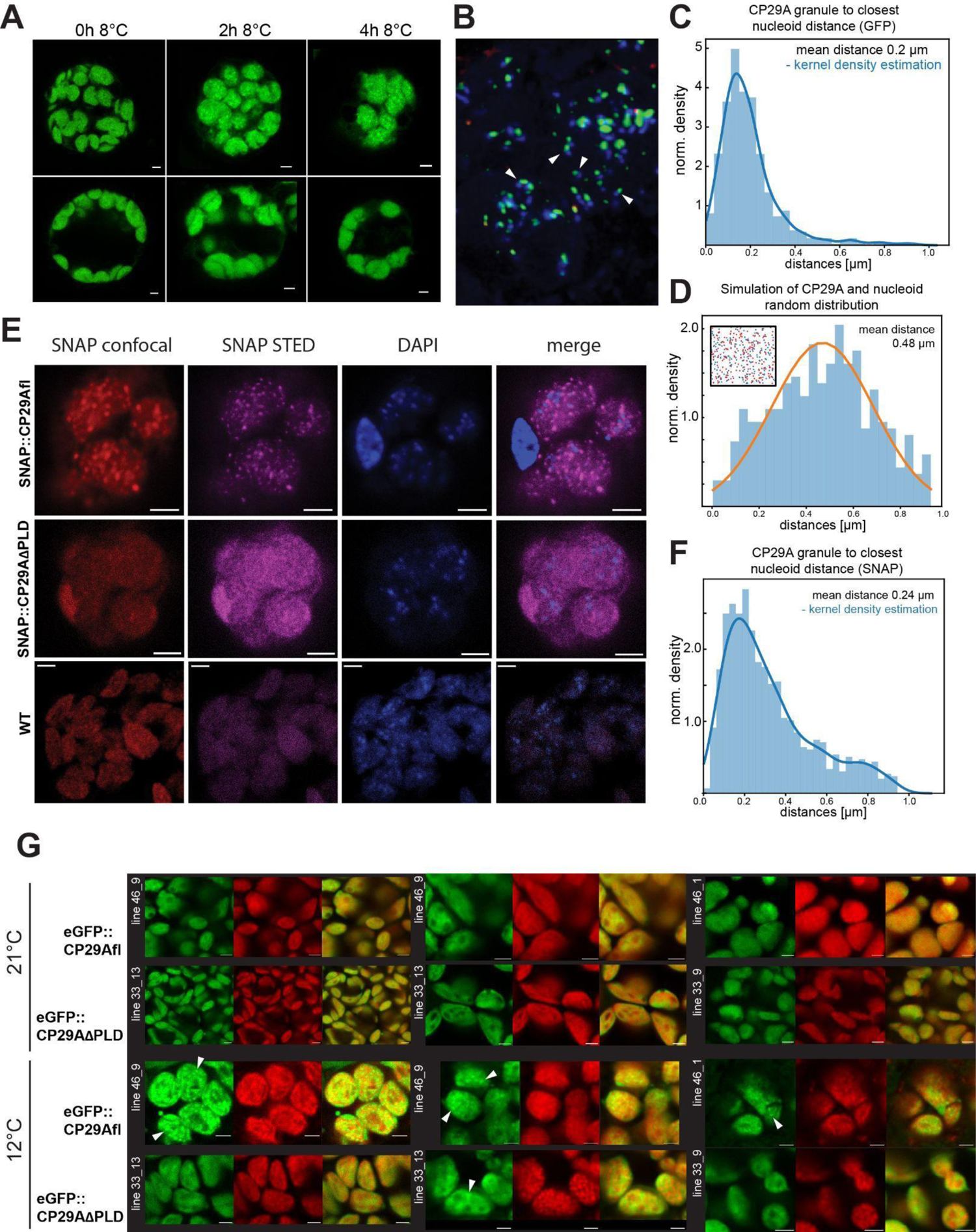
CP29A forms droplets in the vicinity of nucleoids, which depends on its PLD and is fostered by low temperatures (A) Confocal microscopy of protoplasts prepared from wt and the transgenic lines stably expressing mGFP::CP29Afl or mGFP::CP29AΔPLD, exposed to low temperature for the indicated time. Scale bar: 10 µm. Three replicates were performed with independent transgenic lines for each genotype. (B) STED microscopy of protoplasts expressing full-length CP29A fused to GFP and stained with DAPI. Arrowheads point to examples of doublet signals. (C) Quantification of the distance of CP29A droplets and nucleoids in (B). (D) Simulation of a random distribution of CP29A droplets and nucleoids based on the number of signals in (B) per volume analyzed. (E) Confocal and STED microscopy of protoplasts prepared from transgenic lines stably expressing SNAP::CP29Afl or SNAP::CP29A ΔPLD and exposed to low temperatures for different time. Scale bar: 10 µm (F) Quantification of the distance of CP29A droplets and nucleoids in (E). (G) Subcellular localization of mGFP::CP29Afl or mGFP::CP29AΔPLD at standard growth temperatures and after cold acclimation. Two different lines were analyzed for each construct (line numbers in white font). Exemplary granules are indicated by arrowheads. Scale bar: 2 µm

We next investigated whether the formation of CP29A droplets is linked to nucleoids, given that nucleoids contain a large number of RNA binding proteins (Majeran et al. 2011). To answer this question, we utilized STED microscopy, which is better equipped to resolve the suborganellar distribution of CP29A droplets and DAPI-stained nucleoids than standard confocal microscopy. Our observations indicated that the GFP signal and the DAPI signal only occasionally overlap (Figure 3B). However, we frequently observed that the two signals appeared as doublets (Figure 3B arrowheads), prompting us to explore the possibility of a physical connection between CP29A droplets and nucleoids, similar to the known tethering of mitochondrial RNA droplets to nucleoids (Rey et al., 2020; Antonicka and Shoubridge, 2015). By measuring the distance between nucleoids and CP29A droplets, we discovered a non-random distribution of the two signals, with an average distance of 0.2 µm (Figure 3C). As a control, we examined whether a modeled random distribution of nucleoids and CP29A droplets would result in a similar distance. For that we estimated average numbers of nucleoids and CP29A droplets from fluorescence images and found that a random distribution of the two signals in the chloroplast volume led to an average distance of 0,488 µm, which was significantly larger than the distance measured in the in vivo data (Figure 3D).

To validate our findings using an alternative detection method of CP29A droplets, we employed the SNAP tag, which is also more suitable for STED microscopy since it allows the usage of organic dyes that emit two orders of magnitude more photons than GFP (Dempsey et al., 2011; Fernández-Suárez and Ting, 2008) We generated plants that stably express CP29A with an N-terminal SNAP-tag in a *cp29a* mutant background, as well as plants expressing SNAP::CP29AΔPLD in the same background. The expression levels were comparable between both lines and slightly higher than in wt (Figure S3B). We generated protoplasts from both SNAP::CP29Afl and SNAP::CP29AΔPLD lines and incubated them with the SNAP-compatible dye Cell-SiR647 for one hour. As seen in the mGFP::CP29A expressing plants, only the lines expressing full-length CP29A exhibited a granular localization of CP29A, while the plants lacking the PLD showed a diffuse signal with few granular bodies (Figure 3E). Consistent with our observations in GFP-tagged lines, we also found a correlation between the location of CP29A signal and nucleoids in SNAP::CP29Afl protoplasts (Figure 3F), with an average distance of the two signals of 0.24 µm. These findings demonstrate that droplet formation occurs in living chloroplasts and is not reliant on the tags utilized, as both GFP- and SNAP-tagged CP29A form droplets. Moreover, our results provide compelling evidence that CP29A droplets are in close proximity to nucleoids.

To further corroborate our findings, we performed immunofluorescence assays on wild type (wt) protoplasts using an antibody against the native CP29A protein. Since the antibody was designed to recognize the PLD region, it could only be used for analysis of plants expressing full-length CP29A. Consistent with our GFP- and SNAP-tagged fusion protein experiments, immunofluorescence assays revealed granular signals that increased in number at low temperatures (Figure S4). We observed that the granular signals were mainly located near the chloroplast envelope, potentially due to limitations in the antibody’s ability to penetrate formaldehyde-fixed chloroplasts densely packed with thylakoid membranes. Importantly, this analysis demonstrates that droplet formation of CP29A is not an artifact of transgene overexpression or fusion protein production.

To exclude that granule formation is induced by protoplastation, we next investigated leaf tissue for the localization of mGFP::CP29Afl and mGFP::CP29AΔPLD. Starting with material grown entirely at 21°C, we saw very few granular structures in chloroplasts of either genotype (Figure 3G). When the plant material was however transferred to 12°C, the signals became more punctate in mGFP::CP29Afl plants. These signals were no in all cases roundish, possibly restricted and deformed by the thylakoid membrane system. In some mGFP::CP29Afl cells, chloroplasts were replete with granules. Structures with higher signal density were also found in chloroplasts from mGFP::CP29AΔPLD leaves, but in much lower numbers (Figure 3H). Taken together, our analyses provide strong evidence that CP29A forms droplets near nucleoids in living cells, and that this behavior is strongly supported by the PLD region and is promoted by low temperatures.

### The PLD is required for plant cold resistance

Null mutants of CP29A show pale tissue at the center of the leaf rosette after longer cold exposure. We next tested whether the PLD is required for CP29A-mediated cold resistance. To this end, we exposed 14 days old plants, expressing mGFP::CP29AΔPLD and mGFP::CP29A in the *cp29a* mutant background, to 8°C for 14 days. Additionally, we also exposed 14 days old *cp29a*/SNAP::CP29AΔPLD and *cp29a/*SNAP::CP29A plants to the same cold conditions. The plants expressing full-length CP29A remained green under these conditions, no matter which fluorescent tag was used, demonstrating that the N-terminal fusion protein could functionally replace the wt protein (Figure 4A). By contrast, plants expressing PLD-less protein versions showed a pale center of the rosette, although the affected area appeared smaller than in null mutants and the extent of paleness varied between different individual plants (Figure 4A). This demonstrates that the CP29AΔPLD is not capable of fully reverting the mutant phenotype and only leads to a partial complementation. We conclude that the PLD of CP29A is required for cold-resistance, presumably *via* its ability to phase- separate.

**Figure 4:**
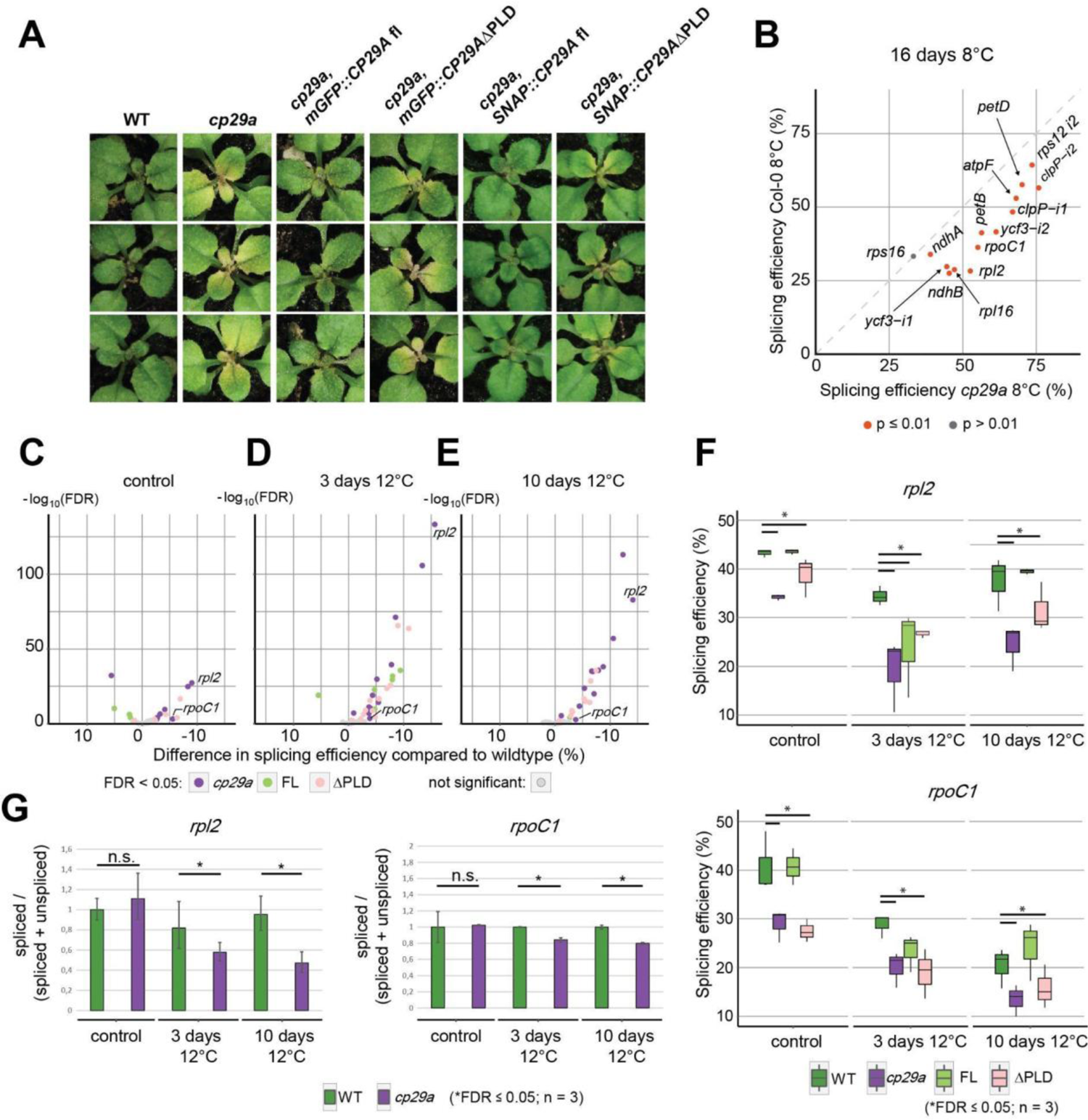
Splicing of chloroplast group II introns in the cold depends on CP29A’s PLD. (A) Phenotype of wt, *cp29a* mutants and complementation lines after 14 days at 21°C and 14 days at 8°C. (B) Analysis of splicing efficiency of chloroplast introns in long-term cold-treated wt and *cp29a* mutant plants by RNA-seq. Splicing efficiency was calculated as the ratio of reads spanning exon-exon junctions versus reads spanning intron-exon boundaries (Castandet et al., 2016). (C) Analysis of splicing efficiency by RNA-Seq after plant growth for 12 days under standard growth conditions in *cp29a* null mutants, wt and complementation lines. Coloured dots represent introns with significantly changed RNA splicing in the different genotypes (FDR > 5). (D) Same analysis as in (C) after 12 days at normal temperature and three days of cold treatment at 12°C. (E) Same analysis as in (C) after 11 days at normal temperature and ten days at 12 °C. (F) Analysis of RNA-Seq data described in (C-E) specifically for the splicing efficiency of *rpl2* and *rpoC1* mRNAs. (G) Analysis of splicing of *rpl2* and *rpoC1* in wt and *cp29a* mutants after three and ten days at 12°C by qRT-PCR. PCR products were either spanning exon-exon junctions (“spliced”) or exon-intron junctions (“unspliced”) and splicing efficiency was calculated as the ratio of qPCR-signals of spliced amplicons over the sum of signal for spliced and unspliced amplicons.

### Chloroplast RNA splicing defects in long-term cold-treated *cp29a* mutants

To understand the molecular function of CP29A in the cold, we initially sequenced RNA from long-term cold-treated mutant plants and compared it to wt plants to get transcriptome-wide insights into potential gene expression defects. We used young plants, 5 days after germination, since the macroscopic phenotype is restricted to young, emerging tissue. These plants were then kept at 8°C for 16 days, during which time the plants hardly grew. Since the chlorotic, cold-dependent phenotypic defect is present in newly developing leaf tissue only, the non-bleached green cotyledons were removed prior to sampling triplicates. RNA was extracted and ribosomal RNAs were depleted prior to Illumina-based sequencing. A principal component analysis was used to ensure the concordance between replicates and experimental groups (Figure S5A). The comparison of the wt plants grown at low temperatures versus *cp29a* mutant plants grown at low temperatures, showed 661 differentially expressed nuclear genes (DEGs), most of them down-regulated in the mutant (Figure S5B). Among the downregulated genes were many photosynthesis-associated nuclear genes (PhANGs). PhANG expression is specifically downregulated at 21°C versus 8°C in mutants, but not in wt, a case in point being the genes for subunits of the light harvesting complexes (LHCs) and photosystems (Figure S5B,C). This is unsurprising, since defects in chloroplast biogenesis are known to induce the reduction of PhANG expression to avoid wasteful protein production when there are problems with chloroplast development (Richter et al., 2023; Liebers et al., 2022). Or in other words, the effects we observed can be explained by the strong chloroplast developmental phenotype and are thus very likely secondary. We noticed, however, an interesting change in splicing efficiency of chloroplast introns in the RNA-seq dataset. The Arabidopsis chloroplast genome contains 21 introns and we tested their splicing efficiency by comparing reads across exon-exon borders versus reads crossing intron-exon borders (introns in tRNAs could not be assessed because their short exons did not allow the detection of intron-exon spanning reads). We obtained splicing efficiency measurements for 15 introns. With the exception of the intron in *rps16,* which is a pseudogene in Arabidopsis and not spliced (Ueda et al., 2008; Roy et al., 2010), splicing was significantly reduced for all introns in cold-treated *cp29a* mutants (Figure 4B).

### Short-term cold-treated plants do neither display chloroplast development defects nor differential chloroplast RNA accumulation

To investigate the onset of molecular defects in the *cp29a* mutant after cold treatment and to avoid measuring secondary effects due to leaf bleaching, we analyzed plants that were exposed to low temperatures for shorter periods. To this end, we harvested tissue that had been grown for 14 days at 21°C and then moved to 8°C for three and ten days, respectively. After three and ten days in the cold, we observed no macroscopic differences between the *cp29a* mutant plants, wt plants and the complementation lines (Figure S6A). To assess the potential occurrence of electron transport problems in the thylakoid membrane after cold treatment, we assayed chlorophyll fluorescence using pulse amplitude modulated fluorescence measurements (PAM). The PAM technique essentially measures changes in PSII photochemical efficiency (in the dark-adapted state), which provides an indicator of electron transport problems in the thylakoid membrane. In long-term treated tissue, *cp29a* mutant plants exhibit a marked increase in fluorescence at the center of the rosette, indicating that normal electron transport in the thylakoid membrane is disturbed (Figure S6B). However, this phenotype is absent in plants exposed to cold for three days, and only a slight increase in fluorescence is measured in *cp29a* plants that were cold-treated for 10 days (Figure S6A).

We then harvested the three youngest leaves, extracted total RNA, depleted rRNA, and performed RNA sequencing in biological triplicates. The three biological replicates showed consistent clustering for the different treatments in principal component analysis (PCA), while the four genotypes showed the least differences under control conditions and the most differences after 10 days of cold treatment (Figure S6C). This supports the idea that expression differences between *cp29a* mutants and wt plants occur only in the cold and increase over time. One replicate of the *cp29a* mutant after 10 days in the cold and one wt plant after three days in the cold seemed closer to the other replicates. However, it is important to note that a greater difference in biological characteristics between samples can occur due to either uncontrolled environmental factors or inherent variations in plant growth.

To investigate potential defects in chloroplast gene expression, we compared the wt, mutant, and complemented mutant samples exposed to the same low temperature regime. Firstly, we looked for differences in chloroplast RNA accumulation, but found none in this short-term cold treated material, including mRNAs for components of the photosynthetic machinery (Figure S6D). This is in contrast to long-term cold-treated mutant tissue, where many photosynthesis-related mRNAs are reduced relative to wt (Figure S5). This analysis demonstrated that the secondary, pleiotropic effects observed in long-term cold-treated *cp29a* plants have not yet manifested after short-term cold treatment and that altered chloroplast RNA accumulation is not an early effect in the mutant in the cold.

### Splicing in short-term cold-treated tissue depends on CP29A’s PLD

To investigate splicing defects in short-term cold-treated plants, we analyzed splicing efficiency in our RNA-seq dataset. Even without cold treatment, there were already several introns with significantly reduced splicing efficiencies in the null mutants and ΔPLD plants relative to wt (Figure 4C). No reduction in splicing was observed in the full-length complementation line, but three introns showed increased splicing efficiency. After three days in the cold, the number of introns with significantly reduced splicing efficiency increased in null mutants from seven to twelve and in ΔPLD complementation lines from five to twelve (Figure 4D). Not only did the number of deficiently spliced introns increase, but the strength of their defects also increased in the cold compared to normal temperatures. Unexpectedly, the full-length complementation line exhibited splicing defects after three days in the cold, albeit to a lesser extent than the null mutant and the ΔPLD line. However, these defects disappeared after ten days in the cold, when only one intron showed significantly reduced splicing in full-length complementation lines (Figure 4E). Given that the full-length construct nearly fully complements the null mutation after ten days in the cold, we speculate that the changes in splicing seen at normal temperatures and at three days cold exposure are caused by expression differences of the transgene relative to the endogenous *CP29A* gene. The genomic location of the construct cannot be controlled by the method used, and thus, despite using the same native promoter for all transgenic lines, different expression patterns can occur.

The PLD-less version of CP29A, in contrast to the full-length construct, still exhibited pronounced splicing defects after ten days of cold treatment (Figure 4E). The defects were not as strong as in the null mutant, indicating that partial complementation of the splicing defect occurs. These findings were also visualized by comparing splicing efficiency of individual introns. For this, we chose *rpl2,* because it showed the strongest reduction in splicing efficiency in the *cp29a* null mutant (Figure 4D,E). *Rpl2* encodes a subunit of the chloroplast ribosome, which is essential for chloroplast development (Zoschke and Bock, 2018). We also added *rpoC1* to the analysis since its gene product is part of an essential chloroplast gene expression machine, the plastid-encoded RNA polymerase (Hajdukiewicz et al., 1997), and is an example of a gene with a milder splicing deficiency in our RNA-Seq experiment (Figure 4D,E). For both introns, the ΔPLD line can only partially complement the splicing defect of the null mutant (Figure 4F). We validated these RNA-seq results by analyzing splicing of *rpl2* and *rpoC1* via qRT-PCR. By amplifying spliced versus unspliced mRNAs, we confirmed the splicing defects of both transcripts in the null mutant (Figure 4G). Overall, our analysis demonstrates that shortcomings in chloroplast splicing are early defects in the *cp29a* mutant after cold exposure and that the PLD is required for full splicing capability in the cold.

### Ribosome profiling of *cp29a* mutants suggests a global reduction of chloroplast translation after short-term cold-treatment

Given that several introns with reduced splicing are in genes coding for ribosomal proteins, including *rpl2*, we hypothesized that *cp29a* mutants may have translation defects in chloroplasts after cold exposure. We therefore used ribosome profiling (Ribo-seq) in *cp29a* mutants and wt to assess translation. Wt and *cp29a* null mutants were grown for 18 days at 21°C and then shifted to 12°C for three days (in three biological replicates). Like in the RNA-seq assays discussed above, the rationale behind this short and mild cold treatment was that we avoid pleiotropic effects of the bleaching phenotype after long-term cold-treated tissue. Whole leaf tissue was processed for Ribo-Seq analysis as well as for parallel RNA-Seq. Ribosomal RNA was depleted from all libraries. Ribo-seq reads mapping to the chloroplast displayed the previously described broad size distribution(Chotewutmontri and Barkan, 2016), whereas reads mapping to nuclear genes resulted in the expected narrower size distribution for cytosolic ribosomes, peaking at 29 nt (Figure S7A). The footprints mapped predominantly to coding regions, where they show the expected periodicity and preference for the first reading frame, (Figure S7B). The Ribo-seq and RNA-seq RPKM values showed a high level of consistency between the replicates (Figure S7C,D), with each replicate comprising at least 2 million Ribo-seq reads and 4 million RNA-seq reads mapping to the chloroplast transcriptome (Table S2).

Standard Ribo-seq and RNA-seq analyses usually rely on detecting a limited number of transcript changes in a large set of transcripts, of which the majority remains unchanged between conditions and genotypes. In a small transcriptome like the chloroplast, a general effect on translation is expected to affect the majority, if not all, of the 80 protein-coding transcripts. Standard normalization of individual sequencing libraries for chloroplast sequencing depth would mask any such general effect. To address this, we compared all chloroplast reads in a sample with all nuclear reads across replicates, thus performing a sample-internal normalization against the nuclear ribosome footprints. This analysis revealed a significant global reduction in chloroplast ribosome footprints in cold-treated mutants versus cold-treated wt (Figure 5A). In contrast, a parallel comparison of the RNA-Seq reads from chloroplasts versus nuclear reads did not show a global change in transcript abundance in the mutant (Figure 5B). This is consistent with our RNA-Seq results from short-term cold-treated mutants and complementation lines: There are no differences in chloroplast RNA accumulation after three and ten days of cold between wt and *cp29a* mutants and there are only a handful of nuclear transcript changes in these libraries (Figure S6D). Similarly, there are no changes in chloroplast transcript abundance and few changes in nuclear transcripts between complementation lines using the full-length CP29A or a PLD-less version of the protein after short-term cold-treated plants (Figure S7D). These results suggest that the reduction in translation observed is not caused by a destabilization of chloroplast transcripts. Therefore, we conclude that translation is globally reduced in *cp29a* cold-treated mutants, likely contributing to the defects in chloroplast biogenesis observed after long-term cold-treatment.

**Figure 5:**
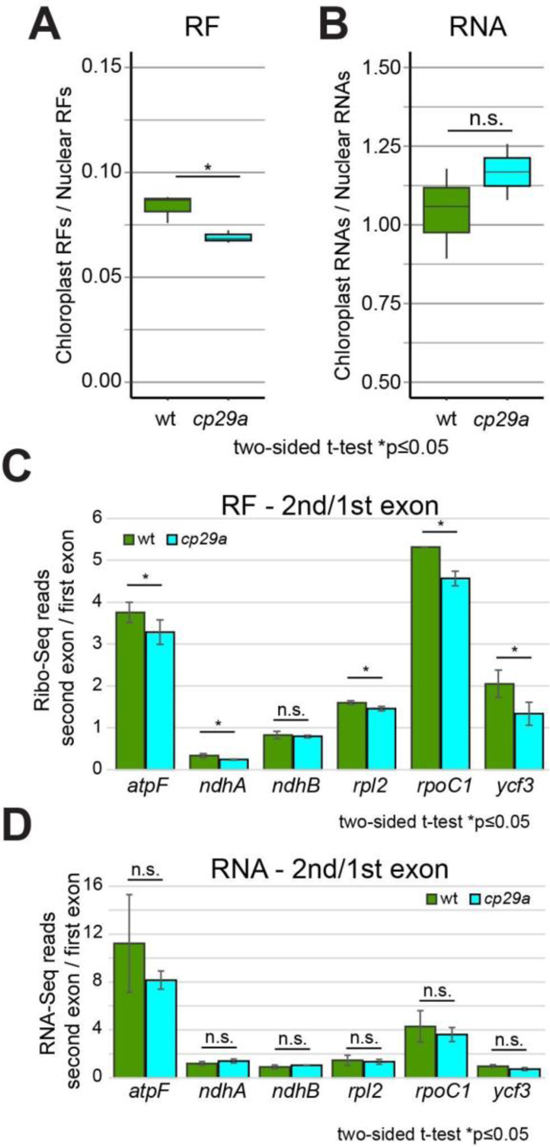
Ribosome profiling and RNA sequencing of short-term cold-treated mutants versus wt. A) The ratio of ribosome footprints from chloroplast versus nuclear mRNAs is shown for cold-treated (12°C - 3 days) wt and *cp29a* mutant plants. The ratios of three individual biological replicates are shown as black dots. Two sided t-tests; *p = 0.001. B) The ratio of RNA-Seq reads for chloroplast and nuclear mRNAs based on RNA preparations from the same batch of plants as in (A). C) Analysis of splicing efficiency in Ribo-Seq data by calculating the ratio of Ribo-Seq reads in the second over the first exon of the chloroplast mRNAs. Two sided t-tests *p ≤ 0.05. D) Analysis of splicing efficiency in RNA-Seq data analogous to (C).

Ribosome occupancy can serve as a readout for splicing efficiency in chloroplasts since, unlike in the nucleo-cytosolic compartment, translation and splicing are not separated (Zoschke et al., 2013; Zoschke and Bock, 2018). As a result of a splicing defect, ribosomes accumulate in the first exon but cannot reach the second exon in unspliced chloroplast mRNAs. Thus, the second-to-first exon read ratio is informative for splicing efficiency. We tested this for all chloroplast introns that had at least 50 reads per adjacent exon. We found a significant reduction in Ribo-Seq reads for the second-exon-over-first-exon ratio for five out of six intron-containing mRNAs in cold-treated mutants compared to wt samples (Figure 5C). In contrast, RNA-seq data show no significant change in RNA coverage between the second and first exon for the same intron-containing genes (Figure 5D). This further supports the requirement of CP29A for efficient splicing of chloroplast introns early after cold treatment. In summary, Ribo-seq reveals that both splicing and translation are reduced in three-day cold-treated mutants.

## Discussion

### The PLD of CP29A is required for efficient chloroplast RNA splicing in the cold

We have established that the chloroplast RNA binding protein CP29A forms droplet-like structures within chloroplasts, which are more prominent in cold conditions. These droplets rely on the presence of the PLD in CP29A, suggesting that they form through phase separation. This is consistent with what we observe in vitro and based on our NMR analysis of full-length and the PLD-only proteins. Our analyses of *cp29a* mutants and complementation lines have revealed that a first detectable effect after cold exposure is reduced splicing efficiency. Even at normal growth temperatures, we have observed minor but significant splicing issues. While splicing is not completely lost in null mutants and ΔPLD mutants after cold treatment, it is noticeably reduced. Based on our findings, we propose several possible explanations for the splicing defect.

Firstly, CP29A may directly act as a splicing factor for chloroplast introns. These introns belong to bacterial group II introns, which require complex 3D structures for proper splicing (Zhao and Pyle, 2017). Previous studies have shown that low temperatures can hinder group II splicing by disrupting the intron’s internal interactions (Dong et al., 2020). We suspect that kinetic traps stabilize alternative RNA structures, which are not biologically functional and impede splicing. In bacteria, RNA binding proteins can aid correct RNA folding at low-temperatures (Lindquist and Mertens, 2018), and bacterial intron maturases can rescue splicing of bacterial introns in the cold (Dong et al., 2020). CP29A may fulfill a similar role by directly interacting with the intron and acting as an RNA chaperone. Arabidopsis CP29A has been previously found to associate with most chloroplast mRNAs in RIP-chip experiments (Kupsch et al., 2012), but a preference for introns was not reported. Relatives of CP29A in tobacco showed a preference for unspliced RNAs in co-immunoprecipitation experiments (Nakamura et al., 1999), and were shown to directly bind RNA in vitro (Ye and Sugiura, 1992; Li and Sugiura, 1991). Thus a direct role of CP29A acting as an RNA chaperone cannot be ruled out at present, but our finding that CP29A forms condensates also supports an alternative mechanism.

Recent studies provide evidence supporting the functional significance of condensate formation in RNA processing. These studies demonstrate that modifying or eliminating protein domains that facilitate phase separation in RNA-binding proteins (RBPs) can impact mRNA maturation (Wang et al., 2018a; Gueroussov et al., 2017; Ying et al., 2017; Li et al., 2020). This phenomenon has also been observed in splicing. Both RBFOX1 and members of the hnRNP A/D family promote splicing of their target RNAs through condensate formation mediated by intrinsically disordered regions (IDR; Ying et al., 2017; Gueroussov et al., 2017). Similarly, the PLD of the *Arabidopsis thaliana* flowering control locus A (FCA) facilitates the creation of nuclear phase-separated bodies that enhance the polyadenylation of target RNAs (Fang et al., 2019). Phase-separated compartments involved in RNA maturation often exhibit higher concentrations of the associated components compared to the non-condensed phase. Such an increase in the concentration of splicing factors would promote the splicing reaction. Chloroplast introns always depend on sets of splicing factors that have overlapping, but not fully redundant intron specificities (recently reviewed in Lee and Kang, 2020; Wang et al., 2022). Speculatively, these chloroplast RNA splicing factors are recruited to CP29A droplets, thus locally enhancing their concentration in the presence of nascent transcripts from the nucleoid. Future cataloging of the components of the CP29A droplets has the potential to determine whether other known chloroplast splicing factors are enriched in the condensed phase.

### Early splicing and translation defects explain impaired chloroplast biogenesis in cold-treated *cp29a* mutants

The first molecular abnormalities observed in *cp29a* mutants after exposure to cold temperatures are a decrease in splicing efficiency and reduced ribosome occupancy of chloroplast mRNAs. These changes occur before any visible effects on photosynthesis and chloroplast development. Conversely, alterations in RNA steady-states in both the chloroplast and nuclear-cytosolic compartments are insignificant. We propose that the splicing and translation impairments are the underlying causes for the inability to produce normally green tissue in *cp29a* mutants under long-term cold conditions for the following reasons.

Firstly, most of the chloroplast genes containing introns are directly involved in the production of the photosynthetic apparatus or encode essential components of the chloroplast gene expression machinery (de Longevialle et al., 2010; Small et al., 2023). For example, the proteins encoded by intron-containing genes *atpF* and *petB/petD* are vital components of the chloroplast ATP synthase and the cytochrome *b_6_f* complex, respectively (Kroeger et al., 2009; Till et al., 2001). Knocking out any of these three genes results in the loss of photosynthesis, leading to pale tissue (Schwenkert et al., 2007) and the absence of splicing in *atpF* alone leads to pale plants and seedling death (Till et al., 2001). Therefore, a reduction in splicing efficiency for *atpF* and *petB/petD* is expected to strain the assembly of the photosynthetic machinery, which becomes critical in cold temperatures where photosystem assembly is expected to be slower.

In addition to the direct impact on photosynthesis gene expression, cold-treated *cp29a* mutants are expected to face general difficulties in producing the chloroplast gene expression machinery, particularly the ribosome. We identified three chloroplast mRNAs encoding ribosomal proteins L2, S12, L16 that show reduced splicing efficiency. Among them, *rpl2* exhibits the most significant splicing defect in our dataset. Therefore, the most plausible explanation for the observed translation defect in cold-treated *cp29a* mutants is the reduced splicing deficiencies of these three mRNAs and thus a limitation in ribosome production. It is however important to acknowledge that we cannot rule out a more direct role of CP29A in chloroplast translation at this stage. The central question remains: How does the reduced translation contribute to the failure to generate green tissue in cold-treated *cp29a* mutants? Loss of the chloroplast ribosome is embryonic lethal in *Arabidopsis thaliana* (Zoschke and Bock, 2018), and hypomorphic mutations in ribosomal protein genes result in the loss of photosynthesis (Dupouy et al., 2022; Bobik et al., 2019). Therefore, the general reduction in chloroplast translation in cold-treated *cp29a* mutants is expected to contribute to the inability to produce photosynthetic tissue. In bacteria, the production and maintenance of ribosomes themselves require specific proteins induced at low temperatures (Jones et al., 1996; Dammel and Noller, 1995), and de novo assembly of newly synthesized bacterial ribosomes is stimulated during cold shock (Charollais et al., 2004). Interestingly, many mutants affecting chloroplast translation exhibit bleaching phenotypes induced by low temperatures, suggesting that any impairment of chloroplast translation will compromise chloroplast development and thus the production of green tissue (Kusumi et al., 2011; Liu et al., 2010; Gao et al., 2022). In fact, a general decrease in translation caused by splicing defects in ribosomal protein mRNAs will also impact the production of chloroplast ribosomes themselves, amplifying the defect. Together, deficiencies in chloroplast translation are likely at the very least contributing to, if not decisively causing, the failure of chloroplast development in *cp29a* mutants in cold conditions.

### Nucleoids as neighbors of a chloroplast RNA processing compartment

Chloroplast DNA is structured into nucleoids, which consist of multiple chromosomes and associated proteins. Our study reveals that the droplets formed by CP29A are primarily found in close proximity to nucleoids. This observation bears similarity to the formation of RNA droplets in the mitochondria of metazoans, which are also spatially associated with nucleoids (Jourdain et al., 2016; Antonicka et al., 2013). The localization of CP29A droplets near nucleoids suggests a potential functional connection between these two cellular components. Just as RNA granules in mitochondria are believed to play a role in RNA regulation and processing, the proximity of CP29A granules to nucleoids implies a similar involvement in chloroplast RNA-related processes. Nucleoid-associated mitochondrial granules contain newly synthesized mitochondrial RNA and contain various RNA binding proteins and RNases required for RNA maturation (Borowski et al., 2012; Pearce et al., 2017). Currently, the composition of CP29A-containing droplets remains unclear, and its protein interaction partners are unknown. Interestingly, a fraction enriched with nucleoids from maize contained factors involved in mRNA processing, splicing, and editing, including homologs of CP29A (Majeran et al., 2011). This led to the suggestion that RNA processing occurs co-transcriptionally. Our identification of a droplet structure adjacent to nucleoids suggests that further subcompartmentalization of gene expression takes place in chloroplasts, separate from nucleoids. The observed distance of 200-240 nm between nucleoids and CP29A droplets implies the possibility of an RNA bridge formed by nascent transcripts (considering that a 1000-nucleotide (nt) RNA molecule is approximately 300 nm in length). This proximity suggests that nascent transcripts could directly interact with CP29A droplets, facilitating RNA processing and other related molecular events. While splicing may be one function of such granular structures, it remains an exciting question for future studies whether other RNA processing steps occur here or in additional separate compartments. The existence of PLDs in a large number of plant proteins suggests that we can anticipate the discovery of many more phase condensation events, including those occurring within organelles (Chakrabortee et al., 2016).

### CP29A droplets as a compartment for cold acclimation

In metazoans, phase separation processes are used as temperature sensors in various contexts (Li and Fang, 2020). Plants, as sessile organisms, need mechanisms to acclimate to changing temperatures. However, examples of temperature-dependent phase separation in plants are rare. In Arabidopsis, increased temperature leads to PLD-mediated phase separation of EARLY FLOWERING 3 (ELF3), which regulates the circadian clock (Jung et al., 2020). Another example is GUANYLATE BINDING PROTEIN LIKE 3 (GBPL3), which undergoes phase separation through a C-terminal IDR, resulting in the activation of target promoters during pathogen infection - a process impaired at higher temperatures (Huang et al., 2021a). Similarly, higher temperatures lead to the dissolution of phase separation-dependent, phytochrome-containing photobodies, thereby modulating light-responsive gene expression (Chen et al., 2022). CP29A differs from these phase separation events in terms of its specific occurrence and importance at low temperatures. Its ability to autonomously and rapidly form condensates in response to lower temperatures in vitro and to occur in granular structures upon cold exposure in vivo together with its cold-specific function in chloroplast development make it a possible chloroplast cold sensor.

At first glance, CP29A droplets appear similar to cytosolic stress granules in mammals and yeast, which can be induced by low temperatures (Hofmann et al., 2012). However, stress granules mostly repress translation and may eventually contribute to RNA degradation (Hofmann et al., 2012). In contrast, CP29A droplets seem to support the accumulation of spliced transcripts and facilitate translation. We interpret CP29A as a factor that helps maintain the chloroplast gene expression program under low-temperature conditions, while stress granules aid in terminating a gene expression program, paving the way for preparing an alternative program to counter the stress (Ivanov et al., 2019). It should be noted that the temperatures used to induce stress granules in mammals are considered well outside the regular physiological range that occurs in mammalian tissues (Lowering from 37° to 10°; Hofmann et al., 2012), while CP29A becomes relevant during mild temperature changes (21°C to 12°C) that frequently occur throughout the seasons in plant leaves. Thus, cold-dependent phase separation of CP29 occurs at relevant temperatures where the ability to sense and react to dropping temperatures is permanently needed.

Another interesting difference of CP29A phase separation compared to other thermo-sensitive phase separation events may lie in the mechanism of droplet formation. Several other RNA-binding proteins that exhibit temperature-dependent phase separation, such as Fused in Sarcoma (FUS), heterogeneous nuclear ribonucleoprotein A1 (hnRNPA1), or the poly(A)-binding Protein1 (Pab1), utilize their RNA-binding domains next to their IDRs to drive phase separation (Molliex et al., 2015; Burke et al., 2015; Yoshizawa et al., 2018; Riback et al., 2017). For example, in the case of FUS, phase separation is supported by the cold denaturation of its zinc finger domain (Félix et al., 2023). Our in vitro and in vivo deletion analyses suggest that for CP29A, the PLD is the dominant factor in mediating cold-induced phase separation. We hypothesize that the RRMs remain intact and uninvolved during phase separation and fulfill important roles inside the droplet, likely in RNA binding. The fact that the PLD-less version of CP29A was still able to partially complement the macroscopic and splicing phenotype of the null mutant in the cold supports this idea. Identifying CP29A’s RNA and protein partners within the droplet will be a crucial next step in understanding the exact mechanism of its effects on splicing and translation. CP29A has been identified in a proteome of heat-induced chloroplast stress granules together with other RNA binding proteins, including chloroplast RNA splicing and RNA editing factors (Chodasiewicz et al., 2020). These granules were induced by treating Arabidopsis seedlings for 30’ min at 42°C, conditions that led to irreversible aggregation of recombinant CP29A in our hands. Targeted proteomic analyses will have to show whether the cold-induced CP29A granules are related to these heat induced stress granules in protein composition and function.

The biochemical and biophysical conditions within chloroplasts with its light-induced changes in pH and redox state are in many aspects different from the nucleo-cytosolic compartment. Oxidative stress has been shown to induce stress granules in chloroplasts from the green algae *C. reinhardtii,* the only other case of a granule affecting gene expression in chloroplasts (Uniacke and Zerges, 2008). Orthologs of cpRNPs are missing in green algae (Ruwe et al., 2011) and stress granules in *C. reinhardtii* are likely functionally and mechanistically quite different from CP29A-droplets. In order to explore the role of CP29A in temperature acclimation it is probably more promising to tap the diversity of Arabidopsis accessions to uncover whether sequence changes in the PLD modulate its ability to phase separate and thus act as part of a thermosensitive response for different temperature ranges.

## Acknowledgements

We are grateful to I. Passow (HU Berlin) for assistance propagating Arabidopsis and F Reiher (HU Berlin) for support with in vitro phase separation experiments. We acknowledge NMR measurements at the Bavarian NMR Center (BNMRZ) at the Technical University of Munich, Germany. We thank Sam Asami, Gerd Gemmecker and Matthias Brandl for technical support with the NMR facility, and Ralf Stehle for his assistance with in-house SAXS measurements. We acknowledge SBGrid and the NMRbox server for providing access to NMR softwares. We are also thankful to Dr. Gero Schlötel from Aberrior Microscope Facility as well as Dr. Karl-Heinz Körtje from Leica Microsystems for their initial support of the STED microscopy. This research was supported by the DFG via TR175-A02 to CSL, SPP1935 to CSL and MS, and GRK1721, SFB1035 to MS.

## Author Contributions

C.S. and M.S. conceived the study, designed the strategy and supervised the experiments. J.L., B.L., N.K. S.G. and Y.G. performed the experiments and interpreted the data. S.M., W.W., E.K. and R.Z. analyzed datasets and interpreted data. C.S. wrote the manuscript, which was edited by all co-authors.

## Material and Methods

### Bacterial Strains and growth

DH5α *E. coli* (Invitrogen-Thermo Fisher, Hennigsdorf, Germany) was used for vector amplification and passing, while BL21 *E. coli* (Invitrogen-Thermo Fisher, Hennigsdorf, Germany) was used for protein expression. *E. coli* cultures were grown at 37C under constant shaking (180 rpm) in standard lysogeny broth (LB) medium. *Agrobacterium tumefaciens* GC3103 cultures were grown at 28°C under constant shaking (200 rpm) in standard lysogeny broth (LB) medium.

### Plant lines

Null mutants of *CP29A* (*cp29a*-6) were obtained from the SALK Institute (San Diego) and were previously described (Kupsch et al., 2012). For transformation, the *cp29a*-6 line was grown under standard conditions (long day conditions; 120 µE s^-1^m^-2^) in a growth chamber. *Agrobacterium* mediated transformation was carried out by standard floral dip methods.

### Plant growth conditions

Arabidopsis seeds were stratified at 4°C in the dark for 3 days and then grown at 21°C with a light intensity of 120 μmol m^−2^s^−1^. For short-term cold treatment, plants were grown for 14 days at 21°C and then transferred to 12°C or 8°C for 3 or 10 days. To analyze the effects of long-term cold stress on *cp29a* and wt plants, both sets of plants needed to be at the same developmental stage. Due to slower growth under cold stress, cold-stressed plants were cultivated for 21 days to reach a comparable developmental stage as unstressed plants grown for 10 days at 21°C. This involved an initial cultivation at 21°C for 5 days, followed by an additional 16 days at 8°C. Before harvesting, cotyledons were removed since the phenotypic effect of a *CP29A* knockout under cold conditions is primarily observed in newly formed tissues.

### Construction of plant transformation vectors and plant transformation

Transformation vectors for the preparation of plant transgenic lines carrying mGFP and SNAP Tags were generated by HiFi assembly. The promoter region of CP29A was amplified with the primer pair pcp29A+70aa_frw and pcp29A+SP70aa_rev, resulting in a 1454 bp fragment (All oligonucleotides used for cloning can be found in Table S3). A monomeric GFP fragment was obtained using the primer pair mGFP FL CP29A_frw and mGFP FL CP29A_rev, yielding a 722 bp fragment. The full-length genomic sequence of CP29A was amplified with the primer pair CP29A genomic+3UTR_frw and CP29A genomic+3UTR_rev, resulting in a 2234 bp fragment. These three fragments were mixed in equimolar concentration with the pGL1 Vector cut with *Bam*HI and *Hind*III. HiFi Assembly was performed following the manufacturer’s protocol (NEB, Frankfurt, Germany), resulting in the generation of a vector named pGL1-pcp29A::mGFP::CP29AFL.

For an analogous SNAP-tagged version of the full-length protein, the promoter region, SNAP Tag, and genomic fragment of CP29A were amplified with the following primer pairs: pcp29a+70aa_frw and pcp29A+70aa_rev for the promoter region, SNAptag_frw and SNAPtag_rev for the SNAP Tag, and CP29AFL SNAPtag_frw and CP29AFL SNAPTag_rev for the genomic fragment. The size of the SNAP Tag fragment was 579 bp, while the sizes of the other fragments were as described above for the GFP construct. After HiFi assembly into pGL1, the vector was named pGL1-pcp29A::SNAP::CP29AFL.

To obtain a version of the constructs without the PLD domain, additional primer pairs were used. The promoter region and the GFP tag were amplified using the same primer pairs as before. A first genomic CP29A fragment, encompassing the RRM1 region, was amplified with the primer pair CP29Amidpart-PLD_frw and CP29Amidpart-PLD_rev, resulting in a 927 bp fragment. The second part of the genomic CP29A sequence, excluding the PLD domain, was amplified with the primer pair CP29ACterm-PLD_frw and CP29ACterm-PLD_rev, resulting in a 1230 bp fragment. After combining these PCR fragments with the pGL1 Vector, a HiFi assembly reaction was performed, resulting in the generation of a vector named pGL1-pcp29A::mGFP::CP29A DPLD.

After confirming the sequence of the PLD-less mGFP-tagged construct through Sanger sequencing, the primer pair for the full-length SNAP Tag construct was used on the template of a GFP construct lacking the PLD domain. This allowed for the generation of a SNAP Tag version of the construct without the PLD domain, resulting in a PCR fragment of 2139 bp. The resulting vector was named pGL1-pcp29A::SNAP::CP29A DPLD. Primer design was performed using NEBuilder v2.8.1 (https://nebuilder.neb.com/#!/).

Bacteria carrying all fragments in the pGL1 Vector derivatives were selected on LB media supplemented with Kanamycin. Transgenic plants generated by Agrobacterium mediated transformation were selected on soil by spraying with BASTA. At least three independent lines per construct were isolated.

### Photosynthetic measurements

In vivo chlorophyll a fluorescence was monitored with the Imaging PAM chlorophyll fluorimeter (Imaging PAM, M-Series; Walz, Effeltrich, Germany). Plants were dark-adapted for 15 min and measured according to standard protocols (Klughammer and Schreiber, 2008).

### Construction of Bacterial Expression Vectors

Vectors for the expression of the CP29A protein in *E. coli* were prepared using the HiFi Assembly protocol with primer-based cloning. To amplify the full-length version of the CP29A protein, the primer pair Nitin_RRM1+2ext(wt)fw and Nitin_RRM1+2ext(wt)rev was used, resulting in a 837 bp fragment. This fragment was assembled into the pETM11 vector backbone, which had been cut with *Nco*I and *Bam*HI, generating a vector named pETM11-CP29A FL.

Similar protocols were followed for the preparation of constructs for NMR measurements. For the construct carrying only the first RRM1 domain, the primer pair FR09 and FR10 was used, resulting in a 321 bp fragment. The fragment carrying the RRM2 domain was amplified using the primer pair FR11 and FR12, also resulting in a 321 bp fragment. Each of these fragments was inserted into the pETM11 vector, which had been cut with *Nco*I and *Bam*HI, using HiFi Assembly.

For cloning of the PLD-only domain, the primer pair Nitin_PLD_fw and Nitin_PLD_rev was used, resulting in a fragment of an unspecified size. This fragment was cloned into the pTrx Vector backbone, which had been cut with *Nco*I and *Bam*HI, and processed with the HiFi Assembly Kit (NEB, Frankfurt, Germany). The resulting vector was named pTrx-CP29APLD core.

For droplet formation assays, constructs were prepared in the pETM11 vector backbone. These constructs contained the mNeon Green Tag fused to the N-terminal side of the CP29A protein. The CP29A full-length version was amplified using the primer pair pETM11 mNeonGreen frw and mNeonGreen29Arev which resulted in a PCR fragment size of 956bp. The mNeonGreen Tag was amplified separately using the primer pair mNeonGreen29A_frw and pETM11 29A_rev which resulted in the fragment size of 753bp. The combination of these PCR products with the NcoI and BamHI-digested pETM11 vector produced the pETM11-mNG::CP29AFL expression vector.

The PLD domain fused to the mNeon Green Tag was amplified using the primer pairs pETM11-mNeonGreen_frw and mNeonGreen_PLD_cor_rev and the PCR product obtained was of 753bp in size, as well as mNeonGreen PLD_cor_frw and pETM11_PLD_cor_rev which amplified a PCR product of 264 bp in size. The resulting fragments were used for HiFi assembly to generate the pETM11-mNG::PLDcore vector.

### Protoplast Isolation and Microscopy

For protoplast isolation, plants were grown on soil for 14 days at standard growth conditions and used for protoplast isolation (Yoo et al., 2007). 10^6^ protoplasts were used for in vivo labeling. SNAP-tagged proteins were labelled with Cell-SiR647 (NEB, Frankfurt, Germany). Labeling was done at RT for 1hr, according to the manufacturer’s protocol. After labeling, a wash step with MMG Buffer was included before fixation. Protoplasts were fixed in ice cold 4% paraformaldehyde in the PBS buffer pH7.2 for 15 minutes. After fixation, three wash steps with ice cold methanol were used for chlorophyll extraction. After a PBS wash, DAPI staining (50 ng/ml) was done in PBS buffer at RT for 30 min. The DAPI stain was washed off 5 times with PBS Buffer. Slides with fixed protoplasts were analyzed using different microscopes. Fig. 3A: Abberior Facility Line Confocal Microscope (Aberrior GmbH, Göttingen, Germany). Fig. 3B: Protoplasts were mounted in Prolong Mountant (Invitrogen, Hennigsdorf, Germany) for STED Microscopy on a Leica TCS SP8 3X (Leica, Wetzlar, Germany). A deconvoluted image of the STED scan is shown, performed with Leica Las X & Huygens Deconvolution software. Fig. 3E: Protoplasts were mounted in Mowiol (Fang et al., 2019) for STED and confocal imaging using the STED Abberior Facility Line machine (Abberior, Göttingen, Germany) and a 100x oil-immersion objective. The DAPI signal of nucleoids was excited with a 405 nm laser, and the emitted light was collected with a detector in the range of 415 to 525 nm. The SNAP Cell SiR647 dye was excited with a 640 nm laser in confocal or STED mode, and the emitted light was collected within the range of 650 to 735 nm for autofluorescence of chlorophyll and fluorescence of SiR647, respectively. SNAP dye-labeled droplets were detected by STED microscropy using excitation at 640 nm and a detecting spectrum of 650 to 735 nm, while the STED laser emitting light used here was at 775 nm with a gate of 750 ps (+8 ns). Fig. 3G: Confocal images from live cell imaging of plant tissue grown at room temperature and plants exposed to cold conditions for 3 days at 12°C. Simultaneous images were taken in the eGFP and STAR635 channels in order to detect eGFP and chlorophyll autofluorescence. Excitation of EGFP was performed with a laser at 518 nm, and the emitted light was collected in the range of 520 to 550 nm, while the STAR635 signal was excited with a laser at 640 nm, and the emitted light was collected within the range of 645 to 670 nm.

### Immunofluorescence assay of CP29A droplets in protoplasts

Protoplasts were allowed to adhere for 10 minutes at room temperature on poly-L-lysine-coated slides. The protoplasts were then fixed with 4% formaldehyde (w/v) in PBS (137 mM NaCl, 2,7 mM KCl, 10 mM Na2HPO4, 2 mM KH2PO4) for 20 minutes at room temperature. To extract chlorophyll, the cells were incubated for 2 x 10 minutes at −20 °C in pre-cooled methanol. Afterwards, the fixed cells were washed for 2 x 10 minutes in PBS-Mg (5 mM MgCl_2_ in PBS). Permeabilization was carried out at room temperature for 10 minutes in 2% Triton or 4% DMSO in PBS. The protoplasts were then equilibrated for 2 x 10 minutes in PBS-Mg and subsequently used for immunofluorescence staining.

The fixed protoplasts were blocked for 30 minutes at room temperature in 0.5% (v/v) BSA in PBS, pH 7.2. The cells were incubated for 75 minutes at 37°C with 2% Triton in PBS for 10 minutes at RT to improve penetration of the antibody. The cells were then incubated with the primary antibody against CP29A(Kupsch et al., 2012; diluted in 0.5% (v/v) BSA in PBS); diluted in 0.5% (v/v) BSA in PBS (dilutions of 1:50 or 1:500 were used). After washing twice for 10 minutes in PBS-Mg, the cells were incubated with a fluorescein-labeled secondary antibody (diluted in 0.5% (v/v) BSA in PBS) for 45 minutes at room temperature in the dark. The samples were then equilibrated again for 2 x 10 minutes in PBS-Mg. Finally, a drop of the permanent mountant VECTASHIELD® HardSetTM Antifade Mounting Medium (Life Technologies, Carlsbad) was added to cover the cells. To preserve the preparation, the specimen was sealed with nail polish.

Epifluorescence microscopy of immunofluorescence assays was carried out with a IX71 ΔVision Spectris Restorations microscopy system (Olympus, Tokyo), employing the 100x Plan Apo objective (numerical aperture 1.4).

### Protein expression and purification

mNG::CP29AFL, mNG::CP29A ΔPLD, and mNG::PLD were expressed in *E. coli* BL21 at 18°C overnight after induction with 0.1 mM Isopropyl β-D-1-thiogalactopyranoside. Bacterial pellets were resuspended in a resuspension buffer (20mM Na-Phosphate Buffer pH 7.4, 300mM NaCl and 1mg/ml Lysozyme). Cells were lysed by sonication and the lysate was centrifuged at 12,000 g for 15 minutes at 4°C. Supernatants were incubated with pre-washed agarose Ni-NTA beads (Cytiva, Freiburg). Bound protein was washed five times with 5 volumes of the bead bed with increasing NaCl concentration (Wash Buffer 1:10mM Tris pH8, 300mMNaCl + 5mM imidazole; Wash Buffer 2: 10mM Tris pH8, 500mM NaCl + 5mM imidazole; Wash Buffer 3:10mM Tris pH8, 1000mMNaCl; Wash Buffer 4: 10mM Tris pH8, 1500mM NaCl). Before elution, a TEV cleavage step was performed with addition of 1000 U of the enzyme per ml of the suspension. Elution of the bound protein was done in 3 Volumes of the bead bed (150 mM NaCl, 10 mM Tris pH 8, 300 mM Imidazole, 1 mM DTT, 0.5mM EDTA). Eluate was concentrated using centricon tubes (Merck, Darmstadt). Protein concentrations were estimated via Bradford assays using a BSA standard. Protein purity was estimated via PAGE-SDS and subsequent CBB staining of the gel.

### In vitro phase separation assays

In vitro phase separation assays with purified recombinant protein were performed in low salt buffer (10mM Tris pH8.0, 50mM NaCl, 1mM DTT, 0,5mM EDTA, 2,5% Glycerol). Crowding agent was added to 10% as a final concentration through mixing with the microscopy solution (20% PEG 8000 in low salt buffer). After mixing, the sample was applied to an 8 well glass bottom slide (Ibidi), incubated at room temperature for 30 seconds before being imaged on an Abberior Facility line Microscope with Light Box software using 100x oil magnification objective. Droplet formation was followed over time by collecting a series of images using a 488 nm excitation laser (3% power). All droplet assays were repeated at least three times. For cold treatment, the protein solution was incubated on ice for 10 minutes prior to microscopy. ImageJ software was used for all image processing Quantifications of droplet size were done with a custom python script in ImageJ. After smoothing and background subtraction using a rolling ball algorithm, droplets were detected with the “Analyze Particles” plugin. The measured Feret’s diameter - maximum diameter of a round object - was then plotted using R-Studio for different conditions.

### Fluorescence recovery after photobleaching

For fluorescence recovery of photobleaching (FRAP) recordings, droplets observed in vitro were excited with 488 nm laser line on confocal Abberior Facility line microscope by using a 100x oil objective. A region of interest was selected within the condensate and 100% power of STED laser was used for bleaching. Five images were taken prior to bleaching and post-bleaching recovery of fluorescence was recorded for up to 55 sec at a frame interval of 1 sec. Evaluation of the FRAP analysis was done with help of ImageJ and custom written Python scripts. In brief, for each condition at least 13 trajectories were analysed, resulting in the fluorescence intensity within the bleached region. First, trajectories were averaged, calculating the standard deviation represented as shadow for each condition in Fig. 1H. Fluorescence intensity recovery trajectories were then fitted using custom written Python scripts with the formula:

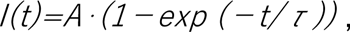

where A corresponds to the plateau intensity and represents the mobile fraction of proteins, τ the recovery time constant and the time after the bleach pulse. Imaging and bleaching conditions such as Laser intensity, bleaching geometry and time were strictly kept constant in between conditions in order to compare mobile fraction and recovery half time (McSwiggen et al., 2019). The value of τ is extracted and is used to calculate the recovery half time t ½:

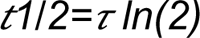

### Distance measurements between CP29A droplets and nucleoids

The shortest distance for every CP29A droplet is measured to its closest nucleoid based on STED microscopy images with an approx. resolution of 60 nm using an Aberior instrument. ImageJ was then used to perform peak detection in both channels. A custom written python script was further used to calculate minimal distances between CP29A and nucleoid pairs. The minimal distances were plotted as a histogram and fitted with a kernel density estimation (KDE). At least 5 field of views containing a minimum of 20 chloroplasts each were analyzed per condition. To confirm the performance of this experiment, we calculated the number of CP29A droplets and nucleoids per chloroplast volume and used a custom written python script to randomly position them. The same script was then used to measure distances. A normalized probability density function to estimate the average distance between centers resulted in a mean of 0.46 um.

### NMR spectroscopy

N-terminal His_6_-tagged RRM1 (97-176 aa), RRM2 (255-334 aa), RRM1,2 (97-334 aa) and His_6_-thioredoxin tagged PLD (176-254 aa) constructs were over-expressed in *E. coli* BL21 (DE3) in M9 minimal media supplemented with ^15^N NH_4_Cl and/or ^13^C glucose as the sole nitrogen and carbon sources, respectively. Cells were harvested, lysed by French press and respective proteins were purified by affinity chromatography with Ni-NTA sepharose followed by TEV cleavage as mentioned above. Further purification steps were carried out by ion - exchange and size-exclusion chromatography. The final buffer for NMR samples contained 20 mM sodium phosphate (pH 6.8), 50 mM NaCl, 1 mM DTT. The sample quality was checked by SDS-PAGE and ESI-mass spectrometry.

Backbone and side-chain chemical shifts for RRM1, PLD and RRM2 constructs of CP29A were assigned from conventional 3D NMR experiments, such as HNCACB, CBCA(CO)NH, HNCO, HN(CA)CO, HNN, (H)CCCONH, H(CCCO)NH and ^15^N-edited NOESY-HSQC (Sattler, 1999; Panchal et al., 2001). NMR measurements were performed on Bruker spectrometers with a proton Larmor frequency of 500, 600, 800, 900, 950 or 1200 MHz equipped with cryogenic or room temperature ^1^H, ^13^C and ^15^N triple resonance probes detection. 5% D_2_O was added to the samples to lock the external magnetic field, NMR samples were put into 3 mm, 5 mm or Shigemi tubes. ^1^H, ^15^N HSQC spectra were acquired with 100 ms and 60 ms acquisition time in direct and indirect dimensions, respectively. Spectra were processed with Bruker Topspin 3.5pl6 or NMRPipe (Delaglio et al., 1995) with a shifted sine-bell window function and zero filling before Fourier transformation. Proton chemical shifts were referenced against sodium 2,2-dimethyl-2-silapentane 5-sulfonate (DSS). Spectra were analyzed by the CCPN (v2.5) software tool (Vranken et al., 2005). Steady-state {^1^H-}-^15^N heteronuclear NOE experiments wererecorded with 170 and 168 ms acquisition times in the direct and indirect dimension, respectively with 3s recycle delay and 32 scans. Spectra were split with a Bruker AU-program, further processed in Topspin and then analyzed using CCPN.

Secondary structural elements for RRM1, PLD and RRM2 domains were derived from NMR backbone and side-chain chemical shifts using TALOS-N (Farrow et al., 1994) and compared with an AlphaFold2 model. The structural model of RRM1-PLD-RRM2 was generated using CNS (v1.2; Shen and Bax, 2013) by keeping the ternary fold of both RRMs (obtained from AlphaFold2), while the coordinates for the PLD linker were randomized. The electrostatic surface charges for CP29a were calculated by APBS tool (Baker et al., 2001).

### Small-angle X-ray scattering (SAXS) measurements

SAXS experiments were carried out in-house on a Rigaku BIOSAXS1000 instrument comprising a a Rigaku HF007 microfocus rotational anode with a copper target (40 kV, 30 mA). Transmissions were measured with a photodiode beam stop and calibration was done with a silver behenate sample (Alpha Aeser). All samples were dialyzed overnight with SAXS buffer (20 mM sodium-phosphate, pH 6.5, 30mM NaCl) before the measurements. Samples with concentrations from 4 to 16 mg/ml were measured at 25°C and 4°C temperature to assess concentration dependent effects. Multiple buffer samples were measured in-between each run and buffer subtraction was applied using the SAXSLab software. Further data analysis was done using the ATSAS package (v3.0.5; Manalastas-Cantos et al., 2021).

### Immunoblots

Equal amounts of total leaf protein were separated on an SDS PAGE gel and blotted to nitrocellulose membrane (Cytiva, Freiburg, Germany). Membranes were blocked in 4% milk TBST solution for 1 hour, washed twice with TBST solution and incubated with a GFP-mouse antisera (Invitrogen, Hennigsdorf, Germany), a CP29A specific rabbit antisera (Kupsch et al., 2012) or anti-SNAP rabbit antisear (NEB, Frankfurt, Germany) in 1:2000 dilution overnight at 4°C and subsequently washed with TBST buffer four times at room temperature. Secondary antibodies (either anti-mouse IgG or anti-rabbit IgG, both conjugated to HRP) were used at a dilution of 1:10.000 for GFP/CP29A primary antisera or at 1:5000 for the SNAP primary antisera and incubated for 1 hour at room temperature. Four washes with TBST buffer were done prior chemiluminescent detection.

### RNA isolation and library construction for RNA-seq

Plant tissue was frozen in liquid nitrogen and pulverized in a mortar. Total RNA was isolated by acid guanidinium thiocyanate-phenol-chloroform-based extraction and purified from the aqueous phase using the Monarch RNA Clean Up Kit (NEB, Frankfurt, Germany). RNA quality was assessed by agarose gel electrophoresis. Total RNA was sequenced on an Illumina platform using Novogene’s (London) lncRNA-seq protocol, based on rRNA-depletion followed by strand-specific library construction. All libraries were sequenced with 150 bp paired-end reads.

### RNA-seq - gene expression analysis

The nf-core/rnaseq (v3.10.1) pipeline was used for quality control, read processing and mapping of the RNA-seq libraries. The Hisat2 was used as aligner and TAIR10 and Araport11 were provided to the pipeline as reference genome and annotation, respectively. FeatureCounts (Rsubread, 2.12.0) was used for read counting and DeSeq2 was used for differential gene expression analysis after low read filtering (CPM>1 in all samples).

### RNA-seq - splicing analysis

Splicing efficiencies were analyzed using the ChloroSeq pipeline, as described previously (Castandet et al., 2016). Fisher’s exact test in combination with Benjamini-Hochberg correction was used for statistical testing.

### qPCR

For qPCR analysis, genomic DNA in total RNA samples was removed using TURBO™ DNase (Thermo Fisher Scientific) followed by purification with the Monarch RNA Clean Up Kit (NEB, Frankfurt, Germany). gDNA removal was assessed by PCR on chloroplast DNA followed by agarose gel electrophoresis. RNA was transcribed to cDNA using the Protoscript®II reverse transcriptase (NEB, Frankfurt, Germany). The Luna qPCR Mastermix (NEB, Frankfurt, Germany) was used for amplification on an 7500 Fast Real-Time PCR System (Applied Biosystems™).

### Ribosome profiling

#### Isolation of ribosome footprints and total RNA

Wt Col-0 and *cp29a-6* plants were grown at 21°C 16/8hrs light/dark regime for 15 days and after this were transferred to cold conditions, 12°C 16/8hrs light/dark regime for three days. 350 mg frozen plant tissue was homogenized in liquid nitrogen with a mortar and pestle followed by the addition of 3.5 mL ribosome extraction buffer (0.2 M sucrose, 0.2 M KCl, 40 mM Tris-OAc pH 8.0, 10 mM MgCl2, 10 mM 2-Mercaptoethanol, 2% (v/v) polyoxyethylene (10) tridecyl ether, 1% (v/v) Triton X-100, 100 μg/mL chloramphenicol, 100 μg/mL cycloheximide). After brief mixing, a 0.5 mL aliquot of the lysate was flash frozen and stored at −80 °C for later total RNA extraction using TRIzol reagent (ThermoFisher cat# 15596026). The remaining lysate was filtered through glass wool, followed by centrifugation for 10 min at 15,000 x g at 4 °C to remove cell debris. 3 mL of the clarified lysate were incubated with 1800 U of RNase I (Ambion cat# AM2294) for 1 h at room temperature with gentle rotation. The ribonuclease-treated lysate contains mainly monosomes which were loaded onto a 2 mL sucrose cushion (30% (w/v) sucrose, 40 mM Tris-Acetate pH 8.0, 100 mM KCl, 15 mM MgCl2, 0.1 mg/mL chloramphenicol, 0.1 mg/mL cycloheximide, 0.2% β-MeEtOH) and centrifuged (OptimaTM L-80 XP Ultracentrifuge - Beckman Coulter) for 1.5 h at 303,800 x g at 4 °C. The supernatant was then aspirated and the pelleted monosomes resuspended in 0.5 mL footprint isolation buffer (10 mM Tris pH 8.0, 1 mM EDTA pH 8.0, 100 mM NaCl, 1% (w/v) SDS, 0.1 M EGTA pH 8.0). The ribosome footprints were extracted from this pellet using 0.5 mL TRIzol reagent.

Next, ribosome footprints were size-selected through electrophoresis on a 12% denaturing polyacrylamide gel (19:1, acrylamide:bisacrylamide) prepared in 1x TBE buffer (89 mM Tris, 89 mM Boric Acid, 2mM EDTA pH 8.0) containing 8 M urea. To this end, Approximately 60 μg ribosome footprints were resuspended in 100 μL of ribosome footprint loading buffer (90% (v/v) deionized formamide, 20 mM Tris-HCl pH 7.5, 20 mM EDTA pH 8.0, 0.04% (w/v) bromophenol blue, and 0.04% (w/v) xylene cyanol) and denatured for 10 min at 70 °C. The gel was run in 1x TBE buffer with a constant power of 30 W at constant temperature of 12 °C (achieved by a cooling unit). Co-migrating pre-stained RNA ladder (Biodynamics Laboratory cat# DM253) was used to visualize the regions of the gel to excise ribosome footprints (20-40 nt). Ribosome footprints were eluted from the excised gel piece in 16 mL TESS (10 mM Tris pH 8.0, 1 mM EDTA pH 8.0, 0.1 M NaCl, 0.2% (w/v) SDS) by overnight incubation at 4 °C with gentle rotation. Eluted ribosome footprints were isolated with 16 mL of phenol:chloroform:isoamyl alcohol (25:24:1), followed by overnight ethanol precipitation at −20 °C. To increase purity and further narrow the volume, the ribosome footprint pellet was resuspended in 0.1 M NaCl (500 μL) and subjected to a second round of phenol:chloroform:isoamyl alcohol (25:24:1) extraction, followed by two washes with chloroform:isoamylalcohol (24:1) and overnight ethanol precipitation at −20 °C. The received ribosome footprint pellet was washed twice with 75% ethanol and resuspended in RNase-free water.

#### rRNA depletion of ribosome footprints

Oligo hybridization was performed following an adapted protocol from (Kraus et al., 2019). 150 ng gel-purified ribosome footprints were mixed with the following components in a PCR tube: 4 μL deionized formamide, 1 μL 20X SSC (3M NaCl, 0.3M Na-Citrate, pH 7.0 with HCl), 2 μL EDTA (5 mM, pH 8.0), 0.7 μL biotinylated oligo mix (100 µM, see table S4), and RNase free water up to 20 μL. Hybridization of the oligos was performed in a thermocycler with heated lid using a slow temperature ramp according to the table below:

After oligo hybridization, each sample was topped off to 40 µL using a solution of 1X SCC and 20% formamide and kept at 35 °C until ready for oligo removal, which was performed using Dynabeads MyOne C1 (Thermo Fisher cat# 65002). For each sample, 45 µL of beads were washed according to the manufacturer’s protocol for RNA applications, and divided into 30 µL and 15 µL aliquots for performing two rounds of oligo removal. Each sample was incubated with the 30 µL bead aliquot at room temperature for 15 min, and magnetized for 2 min. The second round of oligo removal was performed by aspirating the supernatant into the 15 µL bead aliquot and repeating the incubation and magnetization. The final supernatant contains the rRNA-depleted ribosome footprints, which was aspirated, transferred into a clean tube and ethanol precipitated overnight at −20 °C. The rRNA-depleted ribosome footprint pellet was resuspended in RNase-free water and treated with TURBO DNase (Thermo cat# AM2238) according to the manufacturer’s instructions, in order to remove traces of leftover DNA oligos. Following DNase treatment, ribosome footprints were purified using the Monarch RNA cleanup kit (NEB cat# T2030) following the modified protocol for small RNA.

#### Ribo-seq and RNA-seq library construction

To prepare the terminal ends for adapter ligation, rRNA-depleted ribosome footprints were treated with T4 polynucleotide kinase (PNK; ThermoFisher, cat#EK0031). The end-repaired ribosome footprints were purified using the Monarch RNA cleanup kit using the modified protocol for small RNA. These purified ribosome footprints were used as input for the NEXTflex small RNA-seq kit v3 (Perkin Elmer, cat# NOVA-5132-06). The resulting cDNA was amplified by 11-16 cycles of PCR with a barcode incorporated in the primer, and purified according to the instructions of the NEXTflex kit. For RNA-seq, 10 µg of purified total RNAs as described above were subjected to DNase treatment (TURBO DNase, Thermo cat# AM2238) following the manufacturer’s instructions. The concentration of purified total RNA was quantified using a Qubit HS RNA assay (Thermo Fisher Scientific; Q32852). Then, 500 ng of purified total RNA were used to construct sequencing libraries using the Zymo-Seq RiboFree total RNA library kit (Zymo cat# R3000/R3003), which contains a step of an enzymatic degradation of rRNAs. Libraries were pooled for single-end 100-bp sequencing in a NovaSeq 6000 machine. Bioinformatic analysis was done by using the RiboDoc pipeline (François et al., 2021), which features the riboWaltz package (Lauria et al., 2018) for p-site estimation and quality controls.

## DATA AND CODE AVAILABILITY

All data were analyzed using RStudio and Excel. Statistical tests, p values, number of biological and technical replicates, and number of independent experiments are indicated in the figure legends or main text. All RNA-Seq and Ribo-Seq data are available at SRA, bioproject accession PRJNA981550. Reviewer link prior to publication: https://dataview.ncbi.nlm.nih.gov/object/PRJNA981550?reviewer=1crku8tl9crm8itiiqd 3sl742b NMR data are deposited at BMRB with the ID 52022 and 52025.

## Supplemental Data files

**Figure S1:** Sequence, structure and phase separation analysis of recombinant CP29A; related to **figure 1**

**Figure S2:** Small-angle X-ray scattering analysis of the PLD linker of CP29A; related to **figure 2**

**Figure S3:** Accumulation of CP29A variants in transgenic Arabidopsis lines, related to figure 3

**Figure S4:** Immunofluorescence analysis of CP29A during cold acclimation demonstrates an increase in granular CP29A structures in chloroplasts; related to figure 3

**Figure S5:** Principal component analysis and retrograde signaling defects revealed by RNA-Seq analysis of long-term cold-treated *cp29a* mutants; related to figure 4B

**Figure S6:** RNA-Seq analysis of *cp29a* complementation lines after 3 and 10 days in the cold; related to figure 4C-F

**Figure S7:** Quality check of ribosome profiling data of short-term cold-treated plants; related to figure 5.

**Movie S1:** Droplet formation and droplet fusion of mNG::CP29A in vitro; related to figure 1

**Movie S2:** Demixing of PLD domain of CP29A solution that had been kept in ice cold water and was slowly returned to room temperature; A four-minute film shortened to 16 seconds in time-lapse; related to figure 2

