## Supplemental Figures and Tables for "The prion-like domain of the chloroplast RNA binding protein CP29A is required for cold-induced phase separation next to nucleoids and supports RNA splicing and translation during cold acclimation"

### Supplemental Data files

**Figure S1: Sequence, structure and phase separation analysis of recombinant CP29A; related to figure 1**

**Figure S2: Small-angle X-ray scattering analysis of the PLD linker of CP29A; related to figure 2**

**Figure S3: Accumulation of CP29A variants in transgenic Arabidopsis lines, related to figure 3**

**Figure S4: Immunofluorescence analysis of CP29A during cold acclimation demonstrates an increase in granular CP29A structures in chloroplasts; related to figure 3**

**Figure S5: Principal component analysis and retrograde signaling defects revealed by RNA-Seq analysis of long-term cold-treated *cp29a* mutants; related to figure 4B**

**Figure S6: RNA-Seq analysis of *cp29a* complementation lines after 3 and 10 days in the cold; related to figure 4C-F**

**Figure S7: Quality check of ribosome profiling data of short-term cold-treated plants; related to figure 5.**

**Movie S1: Droplet formation and droplet fusion of mNG::CP29A in vitro; related to figure 1**

**Movie S2: Demixing of PLD domain of CP29A solution that had been kept in ice cold water and was slowly returned to room temperature; A four-minute film shortened to 16 seconds in time-lapse; related to figure 2**

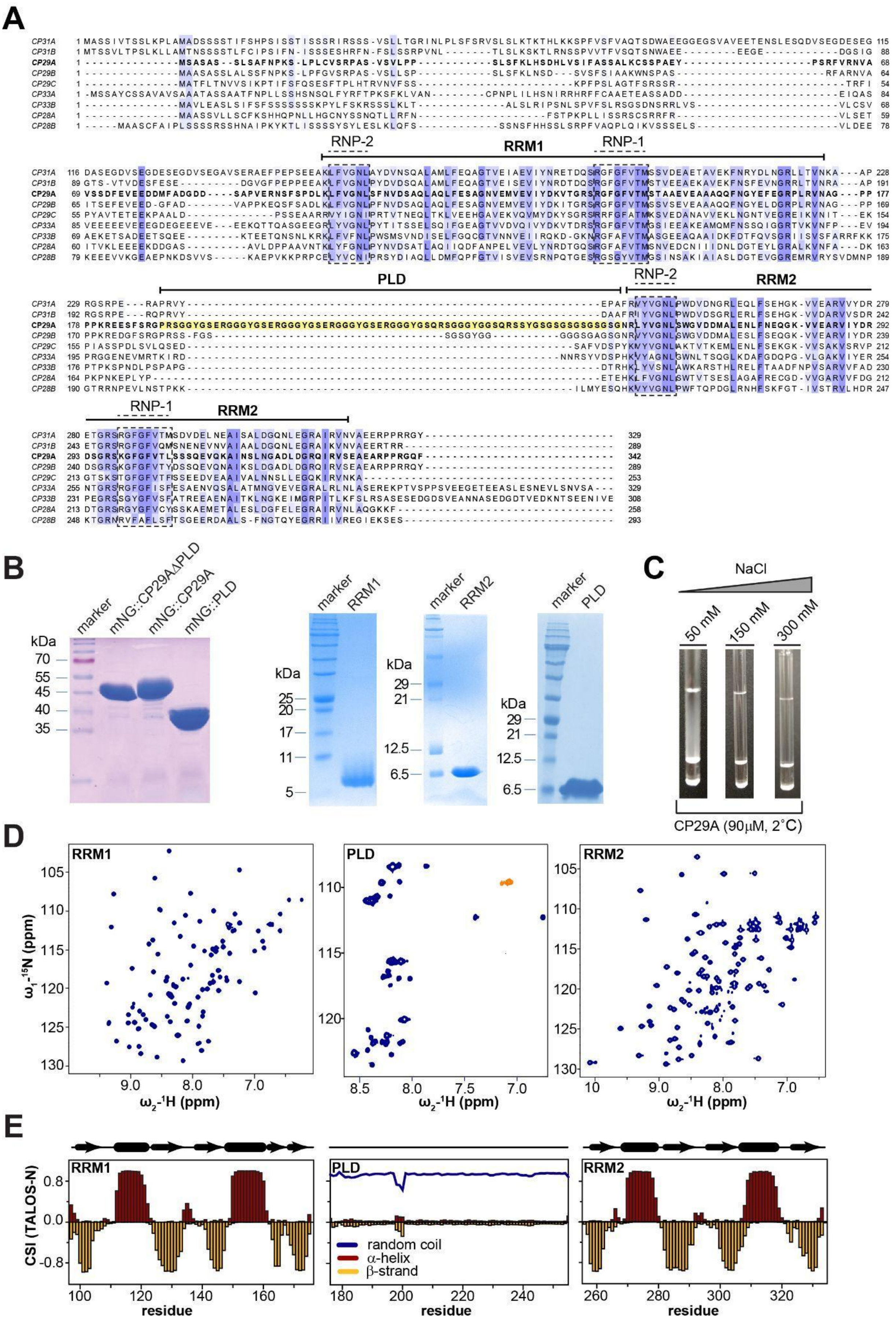


**Figure S1: Sequence, structure and phase separation analysis of recombinant CP29A; related to figure 1**

**(A) Sequence alignment of selected cpRNPs. The PLD of CP29A is highlighted in yellow. Positions of the two RRM motifs and the RNP motifs key for protein-RNA interaction are highlighted by bars.**

**(B) Purified recombinant CP29A protein variants. The RRM domains of CP29A as well as the PLD were expressed separately for NMR analysis. Variants fused to mNeongreen (mNG) were prepared for microscopic analyses. All proteins were expressed in *E. coli* and purified via a $His_6$-tag, which was then removed by proteolytic cleavage. Proteins were separated on a 12% SDS PAGE and stained with Coomassie.**

**(C) CP29A protein solution shows decreased turbidity with increasing salt concentration.**

**(D) $^{1}H,^{15}N$ correlation NMR spectra of RRM1 (left), the PLD linker (middle), and RRM2 (right) of the CP29A protein measured at 298K.**

**(E) TALOS-N derived secondary structure based on NMR backbone chemical shifts for RRM1, PLD and RRM2 of CP29A. RRM1 and RRM2 show the secondary structure expected for a canonical RRM fold, while the PLD domain is unstructured.**

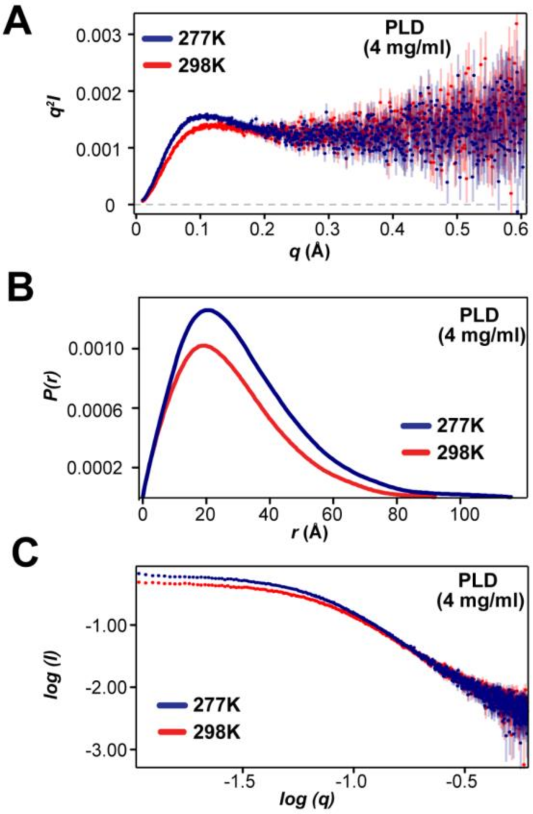


**Figure S2: Small-angle X-ray scattering analysis of the PLD linker of CP29A; related to figure 2**

**(A)** **Experimental SAXS-based Kratky plots of PLD at 4 mg/ml concentration and two different temperatures at 298K and 277K are shown in blue and red, respectively. The plot shape signifies the characteristic of an intrinsically disordered protein.**

**(B)** **The pairwise-distance distribution function *P(r)* of PLD at 298K and 277K is shown in blue and red, respectively, reflecting the overall shape and extension of the PLD linker in solution. At 298K, the maximum distance is ~92Å, while at 274K it is incrased to ~116Å.**

**(C)** **Experimental SAXS data showing the scattering intensity vs. scattering vector at 298K and 277K. The change of intensity at a low scattering angle is consistent with increased molecular interactions.**

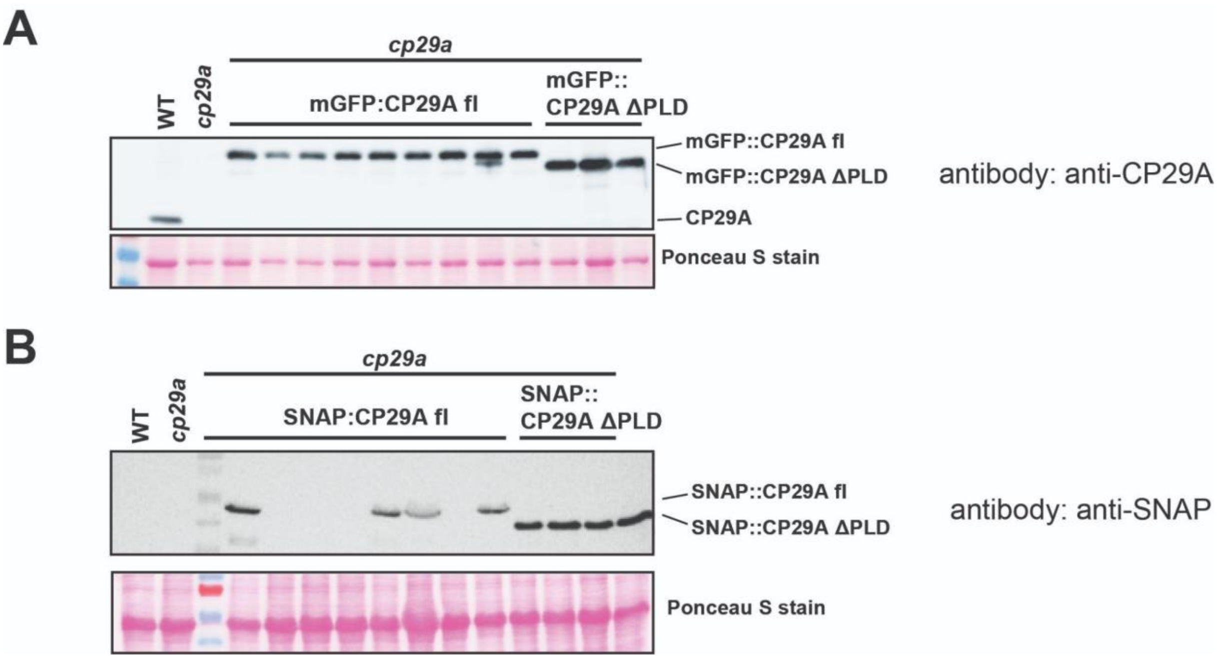


**Figure S3: Accumulation of CP29A variants in transgenic Arabidopsis lines; related to figure 3**

**(A) Immunoblot analysis of mGFP::CP29A fl (full-length) and mGFP::CP29A Δ PLD protein using anti-CP29A antibody.**

**(B) Immunoblot analysis of SNAP::CP29A fl (full-length) and SNAP::CP29A Δ PLD protein using anti-CP29A antibody.**

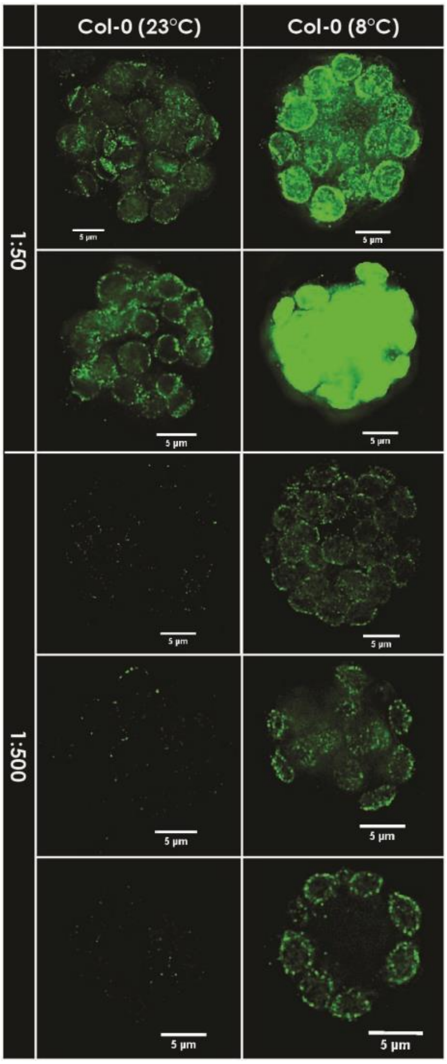


**Figure S4: Immunofluorescence analysis of CP29A during cold acclimation demonstrates an increase in granular CP29A structures in chloroplasts; related to figure 3**

**CP29A fluorescence (FITC, green) is shown for three biological replicates each. Col-0 protoplasts were isolated from four-week-old plants that were either cold-stressed overnight (8°C) or kept at the reference temperature (23°C). Fluorescent labeling was performed with two different concentrations of primary CP29A antibodies (column labels: 1:50, 1:500) and secondary antibodies labeled with fluorescein. All images were made with the same microscope settings, which leads to signal saturation at high antibody concentration in cold-treated protoplasts.**

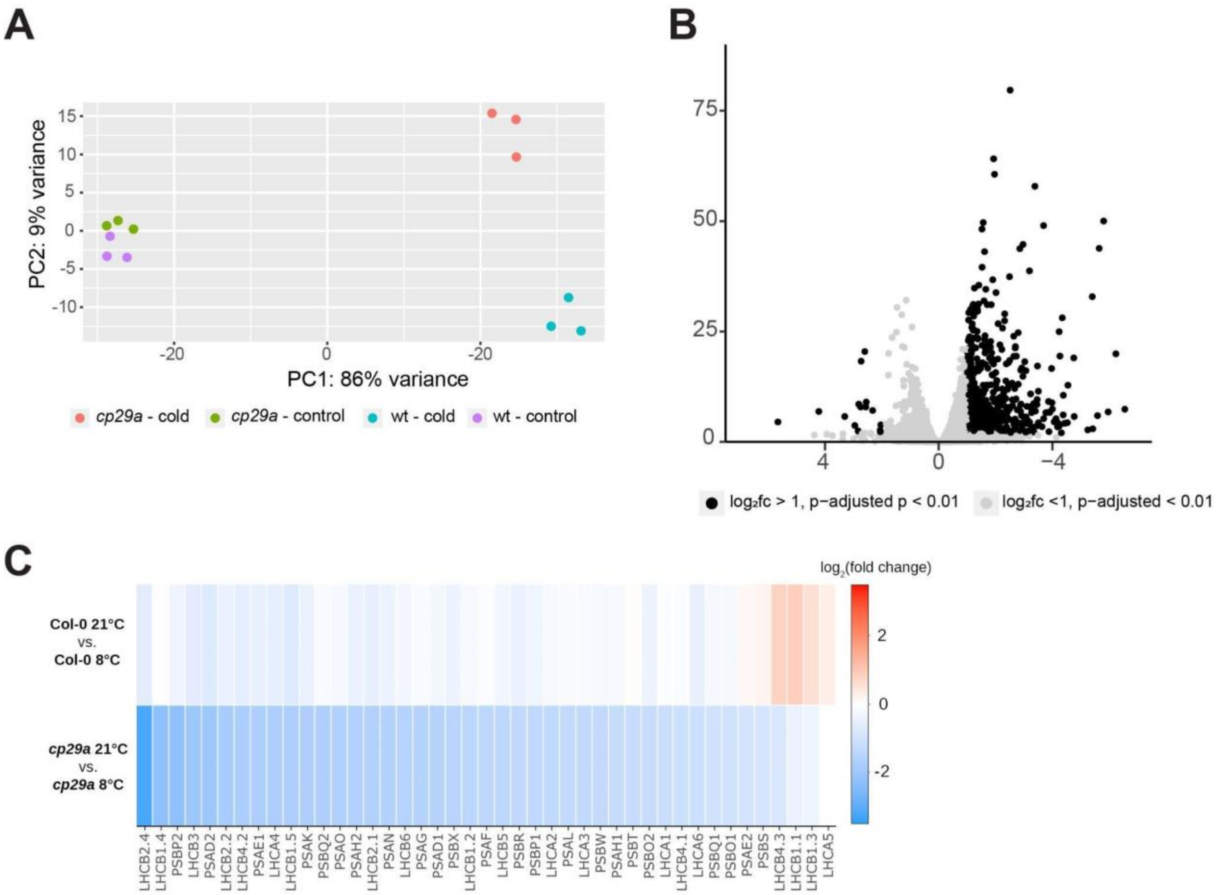


**Figure S5: Principal component analysis and retrograde signaling defects revealed by RNA-Seq analysis of long-term cold-treated *cp29a* mutants; related to figure 4B**

- **(A) Principal component analysis of RNA-Seq data from long-term (16 days) cold treated *cp29a* mutants and wt plants.**
- **(B) Scatter plot summarizing changes in gene expression between long-term cold treated *cp29a* mutants and wt plants. Genes coding for subunits of the light harvesting complexes (LHCs) were marked.**
- **(C) Heat map detailing gene expression changes of genes encoding subunits of the light harvesting machinery of photosynthesis between cold-treated and control plants for PhANGs.**

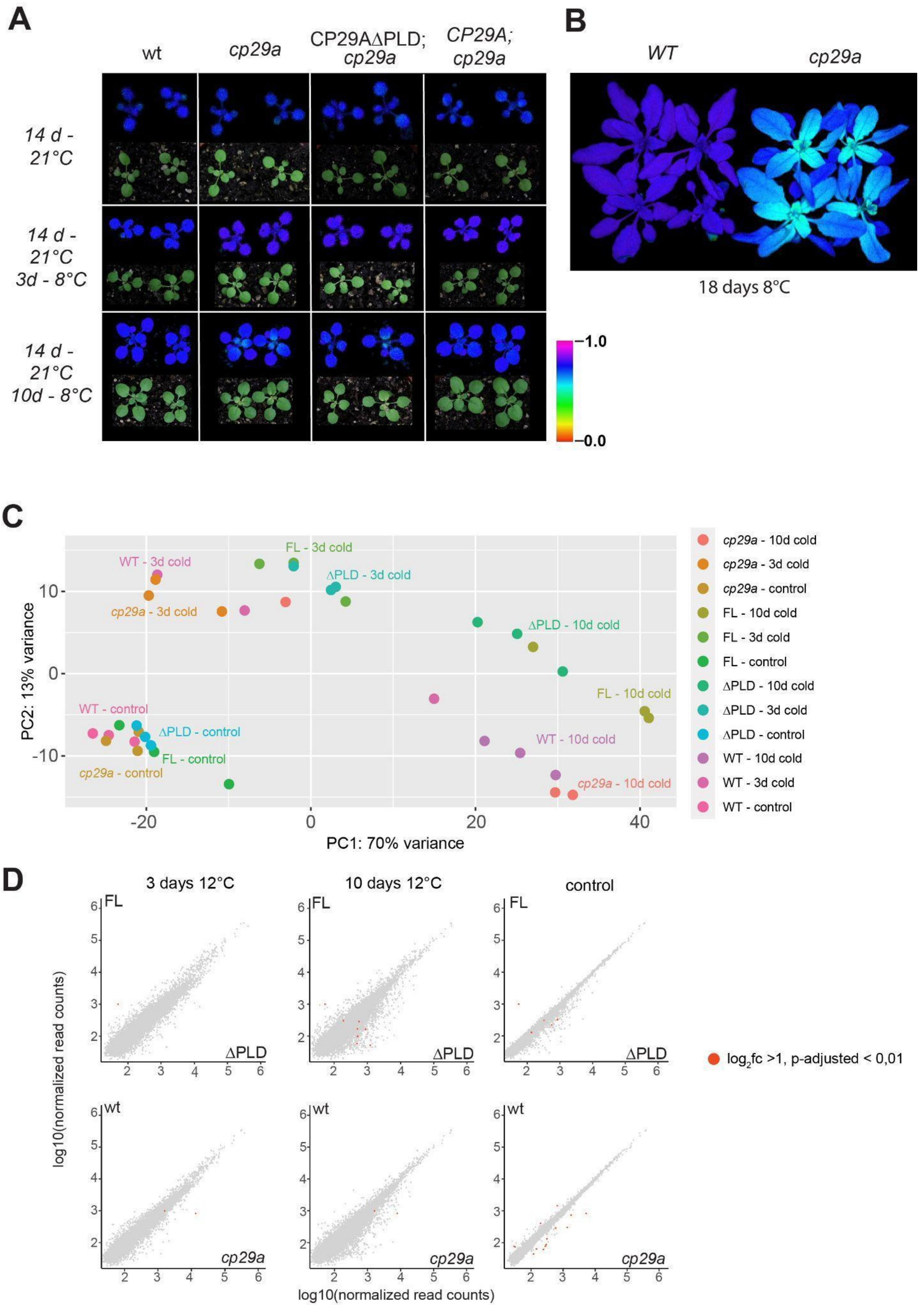


**Figure S6: RNA-Seq analysis of *cp29a* complementation lines after 3 and 10 days in the cold; related to figure 4C-F**

**(A) Macroscopic phenotype and visualization of the maximum quantum yield of photosystem II (FV/FM) with an imaging PAM after three and ten days of cold treatment (8°C). Plants were grown 14 days at standard growth temperatures and then transferred to 8°C for three and 10 days, respectively.**

**(B) Visualization of the maximum quantum yield of photosystem II (FV/FM) after long-term cold-treatment of *cp29a* mutants and wt plants. Plants were grown for 14 days at normal temperatures and then shifted to 8°C for 18 days prior to PAM analysis.**

**(C) Principal component analysis using RNA-Seq reads of short-term cold-treated wt and complementation lines. FL = complementation line expressing the full-length CP29A protein in a null mutant background; ΔPLD = complementation line expressing the PLD-less version of CP29A in a null mutant background.**

**(D) Comparison of RNA-Seq data for complementation lines expressing the full-length protein (FL) or the PLD-less protein (PLD) as well as comparisons of wt and *cp29a* mutants after three and ten days of cold treatment. Comparisons of RNA-Seq results of these genotypes at normal growth temperatures are presented as well (control). Genes with a fold change $\log_2$ >1 and an adjusted $p < 0.01$ are marked in red.**

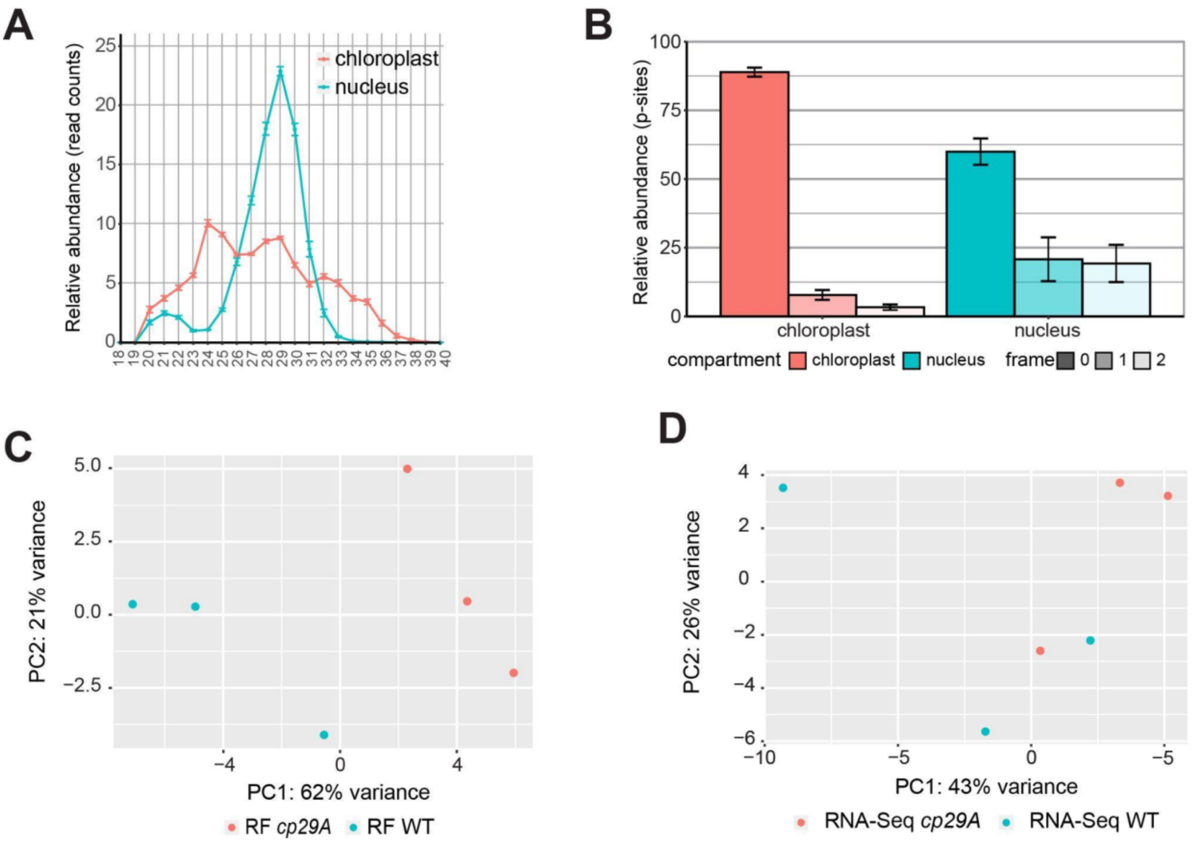


**Figure S7: Quality check of ribosome profiling data of short-term cold-treated plants; related to figure 5.**

- **(A) Size distribution of ribosome footprints. Values are the mean ± SEM from all three samples per genotype analyzed in figure 5.**
- **(B) Three-nucleotide periodicity of Ribo-seq data in CDS versus UTR regions of chloroplast genes. The frame placements were inferred from the locations of the 5' ends of the different footprint sizes at the start and stop codons. Chloroplast footprint in the range of 32-36 nt and nuclear footprints in the range of 29-32 nt were considered for this analysis. Values are the mean ± SEM.**
- **(C) Principal component analysis using Ribo-Seq reads of short-term cold-treated wt and *cp29a* mutants.**
- **(D) Principal component analysis using RNA-Seq reads of short-term cold-treated wt and *cp29a* mutants.**

### Supplemental tables

**Table S1: Small-angle X-ray scattering (SAXS) analysis of the PLD at different temperatures**

| Sample | $R_g$ (Å) | $D_{max}$ (Å) |
|---|---|---|
| PLD (4 mg/ml) at 277 K | 24.24 ± 0.28 | 92 |
| PLD (4 mg/ml) at 298 K | 21.27 ± 0.18 | 116 |

**Table S2: RNA sequencing reads for RNA-Seq and Ribos-Seq experiments of short-term treated wt and *cp29a* mutant plants.**

| | ALL[1] | | NUCLEUS[1] | | CHLOROPLAST[1] | | MITOCHONDRIA[1] | |
|---|---|---|---|---|---|---|---|---|
| | RNA[2] | RFs[3] | RNA[2] | RFs[3] | RNA[2] | RFs[3] | RNA[2] | RFs[3] |
| **wt_1** | 11,809,395 | 24,546,962 | 5,790,067 | 20,920,795 | 5,958,619 | 3,613,177 | 60,709 | 12,990 |
| **wt_2** | 8,811,816 | 14,864,593 | 4,678,168 | 12,792,729 | 4,094,949 | 2,065,365 | 38,699 | 6,499 |
| **wt_3** | 9,625,648 | 25,426,900 | 4,455,407 | 21,712,262 | 5,119,617 | 3,701,006 | 50,624 | 13,632 |
| ***cp29a 1*** | 9,018,417 | 23,563,705 | 4,363,756 | 20,500,187 | 4,605,795 | 3,051,855 | 48,866 | 11,663 |
| ***cp29a 2*** | 14,302,276 | 24,670,729 | 6,648,622 | 21,519,742 | 7,572,629 | 3,135,652 | 81,025 | 15,335 |
| ***cp29a 3*** | 10,154,433 | 22,128,552 | 4,549,498 | 19,280,239 | 5,550,243 | 2,837,669 | 54,692 | 10,644 |

**[1] Numbers for reads mapped to the nuclear, chloroplast and mitochondrial genome of *Arabidopsis thaliana* are shown for each of three biological replicates.**

**[2] Mapped reads for RNA-Seq experiments**

**[3] Mapped reads for ribosome footprints (RFs)**

**Table S3: Oligonucleotides used for cloning**

| ***Name of the oligonucleotide*** | ***Sequence*** |
|---|---|
| pcp29A+70aa_frw | CCCCGGGCTGCAGGTCGACTTGGGACCCATTCTTATTATG |
| pcp29A+SP70aa_rev | TGCTCACCATAGATACAGCAACGTTTCTAAC |
| mGFP_FL_CP29A_frw | TGCTGTATCTATGGTGAGCAAGGGCGAG |
| mGFP_FL_CP29A_rev | CACCTCAAAATCCGACTTGTACAGCTCG |
| CP29A_genomic+3UTR_frw | ACAAGTCGGATTTTGAGGTGGAGGAAGATG |
| CP29A_genomic+3UTR_rev | GAGTGCTTGCGGCAGCGTGAGTTCTACCAATAATATAAATATTCTCTTTTTTC |
| CP29Amidpart-PLD_frw | ACAAGTCGGATTTTGAGGTGGAGGAAGATG |
| CP29Amid_part-PLD_rev | CGTAGAGACGTCCTCTGGAGAAGGATTC |
| CP29A_Cterm-PLD_frw | CTCCAGAGGACGTCTCTACGTGGGCAAC |
| CP29A_Cterm-PLD_rev | GAGTGCTTGCGGCAGCGTGAGTTCTACCAATAATATAAATATTCTCTTTTTTCACC |
| CP29AFL_SNAP_Tag_frw | ATCCCTCGAGGATTTTGAGGTGGAGGAAG |
| CP29AFL_SNAP_Tag_rev | GAGTGCTTGCGGCAGCGTGAGTTCTACCAATAATATAAATATTCTCTTTTTTC |
| SNAPtag_frw | TGCTGTATCTATGGACAAAGACTGCGAAATGAAGC |

| | |
|---|---|
| SNAPtag_rev | CCTCAAAATCCTCGAGGGATCCTGGCGC |
| SNAP_frw_genotype | CAAGCTGGAACTGTCTGGGT |
| SNAP_rev_genotype | CAGGGGATCAGAATGGGCAC |
| pETM11_mNeonGreen_frw | TCCCCACTACTGAGAATCTTTATTTTCAGGGCGCCATGGTATCGAAGGGCGAGGA |
| mNeonGreen29A_rev | TTCTAACGAATTTATACAGTTCATCCATGCCC |
| mNeonGreen_29A_frw | ACTGTATAAATTCGTTAGAAACGTTGCTG |
| pETM11_29A_rev | GTGCGGCCGCAAGCTTGTCGACGGAGCTCGAATTCGGTCATCAAAATTGGCCTCTT G |
| mNeonGreen_PLD_cor_for | ACTGTATAAACCAAGAAGTGGAGGTTAC |
| pETM11_PLD_cor_rev | GTGCGGCCGCAAGCTTGTCGACGGAGCTCGAATTCGGTCATCAGTTTCCTGAACC |
| Nitin_RRM1_plus_2_ext(wt)_fw | TCCCCACTACTGAGAATCTTTATTTTCAGGGCGCCGAGCGTAACTCCTTCTCTCC |
| Nitin_RRM1_plus_2_ext(wt)_rev | GTGCGGCCGCAAGCTTGTCGACGGAGCTCGAATTCGGTCATCAAAATTGGCCTCTT GGTGGC |
| Nitin_PLD_fw | GAGAATCTTTATTTTCAGGGCGCCATGCCTCCTCCACCTAAAAGAG |
| Nitin_PLD_rev | GTCGACGGAGCTCGAATTCGTTATTAACCTGAGCCCGAAC |

**Table S4: Custom biotinylated DNA oligos for rRNA removal from Arabidopsis ribosome footprint samples.**

| ***Name*** | ***Sequence*** | ***Molar Ratio*** |
|---|---|---|
| Nu_5.8S_1 | GGGCGCAACTTGCGTTCAAAGACTCGATGGTTCACGGG | 10 |
| Nu_5.8S_2 | GTGACACCCAGGCAGACGTGCCCTCGGCC | 4 |
| Nu_5.8S_3 | TACGTTCTTCATCGATGCGAGAGCCGAGATATCCGTTGCCGAGAGTCG | 2 |
| Nu_18S_1 | TACCATCAAACAAACTATAACTGATTTAATGAGCCATTCGCAGTTTCACAGTCTG | 2 |
| Nu_18S_2 | TGCACGTATTAGCTCTAGAATTACTACGGTTATCCGAGTA | 16 |
| Nu_18S_3 | CGTCGACCTTTTATCTAATAAATGCGTC | 4 |
| Nu_18S_4 | GGCCATGCGATCCGTCGAGTTATCATGAATCATCAGAGCA | 10 |
| Nu_18S_5 | TCGCCGACCGAAGGGACAAGCCGACCA | 4 |
| Nu_25S_1 | GGGAATCCTTGTTAGTTTCTTTTCCTCCGCTTAT | 2 |
| Nu_25S_2 | CGTCCGATTTTCAAGCTGGGCTCTTCCCGGTTCGCT | 2 |
| Nu_25S_3 | GAGGACGCTTCTCCAGACTACAATTCGAACGCCG | 2 |
| Nu_25S_4 | AGCCCGGGCTTAGGCCGCCACCGTAATCCGCGTCGGTCCACG | 4 |
| Nu_25S_5 | AGGCGCGTGCTGCAGACCACGATCACGGCAGCGACGTCTCCACAAGCGTAT | 8 |
| Nu_25S_6 | AGATCAAGGTCGGTCGGCGGTGCACCCG | 1 |
| Nu_25S_7 | CCGTTCCCAGTCCGTCCCCCGGCCGGCAC | 4 |
| Nu_25S_8 | AGCGAGCCTTGGGACCAAAAACAGGGGT | 4 |
| Nu_25S_9 | GCTCCTACTGAGGGTCGGCAATCGGGCGGCGGGC | 1 |
| Nu_25S_10 | GTCGAATCTTAGCGACAAAGGGCTGAATCTCAGTGGATCGTGGCAGC | 10 |
| Nu_25S_11 | CTCGGTCCTCCGGATTTTCAAGGGCCGCCGGGGGCGCACCGGACACCACGCGA | 2 |
| Cp_4.5S_1 | TAAACGGCTCGTCTCGCCGTGACCTTC | 2 |
| Cp_5S_1 | TGGTGTTGTTCCTCTACGCCTAGGACACCAGAA | 1 |
| Cp_16S_1 | ACGCACAGCGCCTAGTATCCATCGTTT | 1 |
| Cp_16S_2 | TCCCGTCCGACTTGCATGTGTTAAGCATGCCGCCAGCGTTCATCC | 1 |
| Cp_23S_1 | TGCTCTCCCACAACCCCGTTTTC | 2 |
| Cp_23S_2 | GGCTCCTCCCACTGCTTGGGAGCTTACGGTTTCATGTTCT | 1 |
| Cp_23S_3 | GAGGTCATATCTAGTATTCAGAGTTTGCCTCGATTTGGTACCGCT | 8 |
| Cp_23S_4 | AGGTCGTTCGAGCTTTTCCTGGGAGTATAGCATGGGTTACTTCAGCG | 4 |
| Cp_23S_6 | ATCCCACAGCTTCGGCAGATCGCTTAGCCCCGTTCA | 2 |
| Cp_23S_7 | CGCCTGGTACTCGAACATTGGCTCGGGGCATTTTCTCTACCCCTTCTT | 2 |

*Nu, nuclear-encoded; Cp, chloroplast-encoded.*
